# Coming up short: generative network models fail to accurately capture long-range connectivity

**DOI:** 10.1101/2024.11.18.624192

**Authors:** Stuart Oldham, Alex Fornito, Gareth Ball

## Abstract

Generative network models (GNMs) have been proposed to identify the mechanisms/constraints that shape the organisation of the connectome. These models parameterise the formation of inter-regional connections using a trade-off between connection cost and topological complexity or biophysical similarity. Despite their simplicity, GNMs can generate synthetic networks that capture many topological properties of empirical brain networks. However, current models often fail to capture the topography (i.e., spatial embedding) of many such properties, such as the anatomical location of network hubs. In this study, we investigate a diverse array of generative network model formulations and find that none can accurately capture empirical patterns of long-range connectivity. We demonstrate that the spatial embedding of longer-range connections is critical in defining hub locations and that it is precisely these connections that are poorly captured by extant models. We further show how standard measures used for model optimisation and evaluation mask these and other differences between synthetic and empirical brain networks, highlighting the need for care when interpreting generative network models and metrics. Overall, our findings demonstrate common failure modes of GNMs, identify why these models do not fully capture brain network organisation, and suggest ways the field can move forward to address these challenges.

**Author summary:** Generative network models aim to explain the organisation of connectomes using simple wiring rules. While these models replicate topological features of brain networks, they do not capture key topographical properties, like the anatomical location of network hubs. We show that this failure occurs because the models are unable to accurately capture the spatial position of long-range inter-regional connections. Moreover, standard evaluation measures fail to accurately quantify the similarity between model and empirical networks. This study identifies how and why limitations of current generative models occur and suggests ways forward for improved practices.

## Introduction

Dynamic neuronal activity coordinated across distributed brain networks supports a diverse range of complex cognitive processes. This coordination is enabled by the connectome, which comprises the complete array of white matter connections within the brain of any organism (Bullmore & Sporns, 2009). In humans, connectomes are typically studied using non-invasive diffusion magnetic resonance imaging (dMRI), which has been used to reveal a core set of complex topological properties, such as a heterogeneous distribution of connectivity across nodes (Oldham & Fornito, 2019; van den Heuvel & Sporns, 2013), small-world organization (Bassett & Bullmore, 2006, 2017), hierarchical modular architecture (Bassett et al., 2008; Meunier et al., 2009; Sporns & Betzel, 2016) and a rich-club organization in which high-degree hub nodes are strongly interconnected with each other (Arnatkevičiūtė et al., 2021; Oldham & Fornito, 2019; van den Heuvel & Sporns, 2011, 2013).

These topological properties, despite imbuing the network with a certain degree of complexity, may nonetheless arise from simple wiring principles (Albert & Barabási, 2002; Barabási & Albert, 1999; Watts & Strogatz, 1998). These principles can be studied using generative network models (GNMs), which formalize specific wiring rules which are used to construct synthetic networks (Betzel et al., 2016; Betzel & Bassett, 2017; Oldham, Fulcher, et al., 2022; Vértes et al., 2012; Vértes, 2023). Rules that give rise to networks with properties that resemble those observed in empirical data give an indication as to how connectome architecture is shaped. Therefore, generative network modelling provides a useful framework for identifying putative mechanistic or organisational principles that may underlie the formation and development of human brain networks (Betzel & Bassett, 2017; Vértes, 2023). GNMs have also been used to explore how variations in wiring rules relate to individual differences in cognitive performance and psychopathological symptoms (Akarca et al., 2021; Carozza et al., 2023; Vértes et al., 2012; Zhang et al., 2021).

Simple models that attempt to minimise the physical wiring cost of the network (often operationalized as the total connection length of the network) capture many aspects of human connectome organisation (Cherniak et al., 2004; Chklovskii, 2004; Chklovskii et al., 2002; Ercsey-Ravasz et al., 2013; Henderson & Robinson, 2013, 2014; Horvát et al., 2016; Kaiser & Hilgetag, 2004; Raichle & Mintun, 2006; Rivera-Alba et al., 2011, 2014; Song et al., 2014), but often fail to capture properties associated with high-cost, long-range connections (Bullmore & Sporns, 2012; B. L. Chen et al., 2006; Kaiser & Hilgetag, 2006) that act as crucial foundations for dynamic brain function (Betzel & Bassett, 2018; Bullmore & Sporns, 2012; Deco et al., 2021; Kaiser & Hilgetag, 2006; Sato, 2021).

An alternative class of models that attempts to overcome the limitations of cost-only models are based on cost-value trade-offs (Bullmore & Sporns, 2012; Y. Chen et al., 2013; Schröter et al., 2017; van den Heuvel et al., 2016). These models often specify some trade-off between a penalty on long-range connections (i.e., to minimise wiring costs) and a bias towards forming connections that enhance topological complexity in some way (Akarca et al., 2021; Betzel et al., 2016; Carozza et al., 2023; Vértes et al., 2012); for caveats, see Oldham et al., 2022). Such trade-off models typically perform better than those based on cost-minimization alone, particularly when the topological bias favours a form of topological homophily, in which connections are favoured between nodes with similar neighbourhoods (Akarca et al., 2021; Arnatkevičiūtė et al., 2021; Betzel et al., 2016; Y. Chen et al., 2017; Goulas, Betzel, et al., 2019; Vértes et al., 2012; Zhang et al., 2021). Formulations incorporating wiring rules based on the molecular similarity of brain regions have also proven successful (Arnatkevičiūtė et al., 2021; Oldham, Fulcher, et al., 2022), following evidence that cortical regions with similar cytoarchitectural, molecular, and laminar organization are more likely to be connected (Barbas, 1986; Fornito et al., 2019; Hilgetag et al., 2019).

The biological insight offered by GNMs rests on their ability to accurately model empirical human brain networks. While adept at generating networks with similar topologies, GNMs generally fail to capture the way in which these properties are spatially embedded––i.e., the topography of the connectome (Akarca et al., 2021; Arnatkevičiūtė et al., 2021; Y. Chen et al., 2017; Oldham, Fulcher, et al., 2022; Zhang et al., 2021). For instance, the models can often capture the degree distribution of the empirical data, including the existence of hubs, but the hubs reside in very different anatomical locations to those of actual connectomes (Arnatkevičiūtė et al., 2021; Oldham, Fulcher, et al., 2022). This oversight is critical given that the brain’s geometry constrains its function (Pang et al., 2023), and cortical hubs are positioned to achieve a near-optimal trade-off between wiring costs minimization and promoting complex, efficient brain dynamics (Gollo et al., 2018; Roberts et al., 2016). Topographical properties of the brain thus represent important features that generative models of brain networks should capture.

In this study, we perform a comprehensive evaluation of the ability of GNMs to reproduce both the topology and topography of human brain networks. We first demonstrate how and why GNMs fail to capture topographical features of brain networks. Secondly, we examine why the objective functions commonly used to fit GNMs are not sensitive to these differences, finding they can obscure the true extent of differences between networks and are highly sensitive to the presence of certain network properties, but not others. Our findings highlight where GNMs have weaknesses and how the choice of measures used to evaluate them may mask discrepancies with empirical connectomes. By identifying these weaknesses, we outline potential future avenues to address them.

## Results

### Generative models of the human connectome

We defined GNMs using a standard formulation (Betzel et al., 2016; Oldham, Fulcher, et al., 2022). Given a network of n nodes, the models add connections sequentially to populate an empty n × n adjacency matrix, A, according to the wiring-rule

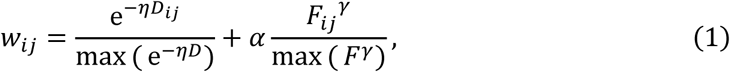

where D_ij_ is the wiring distance between nodes i and j, and F_ij_ is a pairwise interaction for a given feature (either biophysiological similarity or topological coupling). The parameters 1, y and a control the strength of the distance penalty, the scaling of the feature pairwise values, and the relative contribution of the feature term respectively. Each term is normalised by the maximum value of connections which are not already present in the network. The value of w_ij_ is then used to inform the probability of a given connection being generated at each timestep of the model (**Figure 1A,B; Methods;** alternative formulations were explored in supplementary analyses).

**Figure 1.**
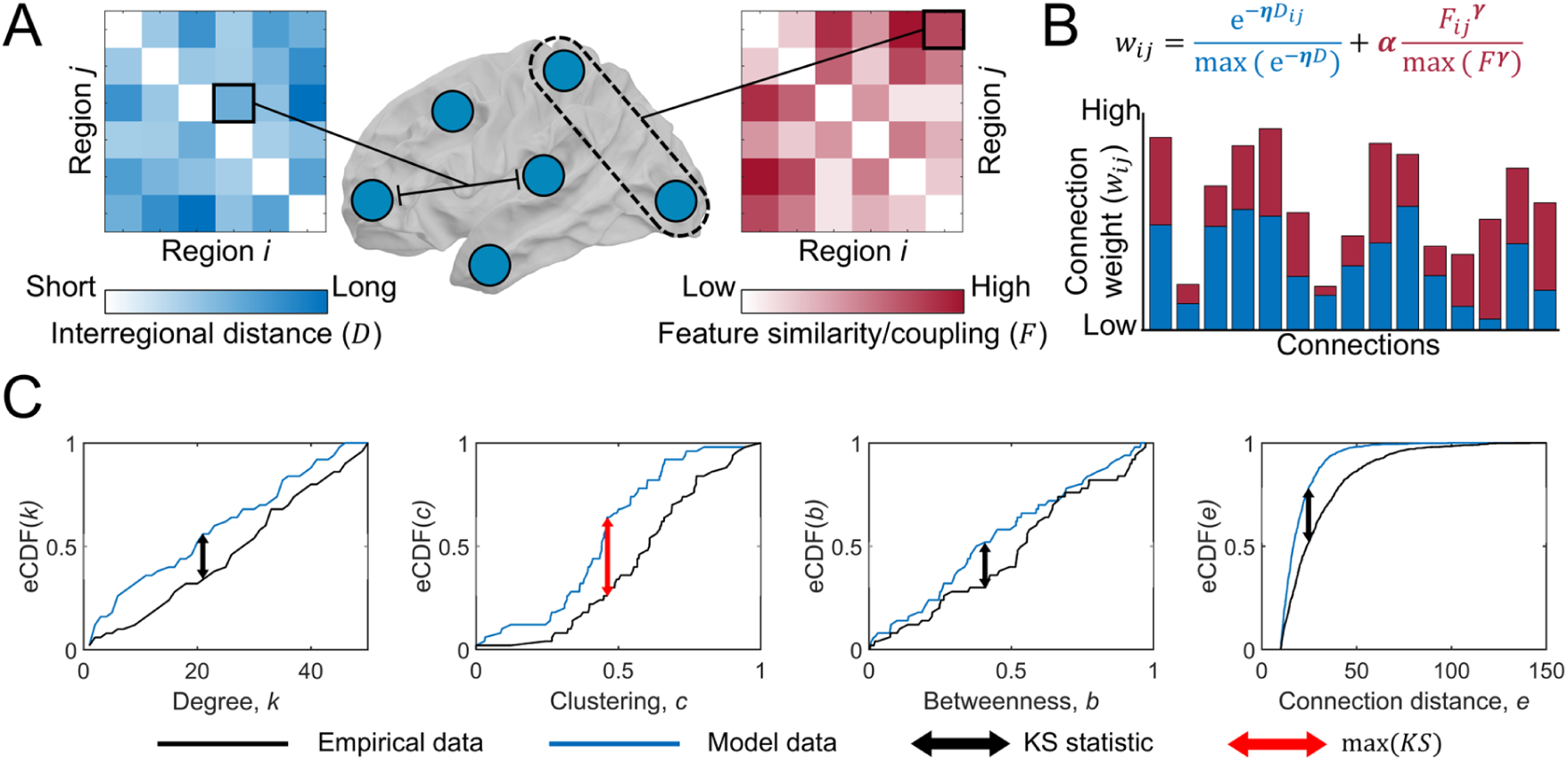
Schematic of the generative network model and max(KS) calculation. **(A)** The distance (wiring cost; D_ij_) and similarity/coupling on a given feature (F_ij_) between nodes i and j is used in the generative network model. F_ij_ can represent either inter-regional similarity on some biophysiological feature, or coupling of some topological feature. **(B)** Each connection is assigned a weight w_ij_ according to the equation shown, each defines a trade-off between the cost of a potential connection (e^-r,Dij^) against the regional similarity F_ij_^y^. Each term is normalised by its respective maximum value. Three parameters 1, y and a control the strength of the distance penalty, the scaling of the feature pairwise values, and the relative contribution of the feature term respectively. The model adds connections on an iterative basis to a network based on w_ij_ (higher values indicate a greater likelihood of forming, bar plot). **(C)** Topological similarity between empirical and synthetic networks is defined as the maximum Kolmogorov–Smirnov (KS) statistic max(KS) across four topological distributions: degree, clustering, betweenness, and connection distance. The arrows indicate where the KS statistic is calculated; the red arrow indicates the maximal difference across the four properties (i.e., max(KS)).

The parameters of the model were formulated to always impose a penalty of long-range connections (the strength of which could vary), while affording the flexibility to form connections between regions with similar or dissimilar nodal properties (Oldham, Fulcher, et al., 2022). The value of F_ij_ could indicate either a measure of pairwise similarity for a cortical feature (e.g., correlation of two regions gene expression patterns), or indicate some level of topological coupling (e.g., the matching index, which quantifies the similarity of the topological neighbourhoods between nodes) between the pair of regions that is continuously updated as the model progressively adds connections to the network.

We defined a total of 10 different GNMs (**Table 1**), which included: one where pairwise similarities between nodes F_ij_ was defined using the matching index (*Matching*); seven with similarity defined by different cortical biophysiological features, which include similarity in gene expression, receptor density, lamination, glucose uptake, haemodynamic activity, electrophysiological activity, and temporal profiles (Hansen et al., 2023); and two baseline models, one using wiring distance only (*Spatial*); and one where nodal similarity was based on random, spatially autocorrelated data (*Random similarity*). We focused on these 10 models as: a) previous work has found that biophysiological similarity GNMs outperform topological GNMs (Arnatkevičiūtė et al., 2021; Oldham, Fulcher, et al., 2022); b) these seven biophysiological measures represent a comprehensive summary of different measurements of inter-regional similarity (Hansen et al., 2023); c) the *Matching* model is often found to be the best performing topological model so was included as a point of comparison; and d) the two baseline models are appropriate nulls given the impact of distance and spatial-autocorrelations in shaping cortical features and connectivity. In supplementary analyses, we examined 12 topological GNMs (**Table S1**) that have been widely used previously (Akarca et al., 2021; Arnatkevičiūtė et al., 2021; Betzel et al., 2016; Oldham, Fulcher, et al., 2022).

**Table 1.** Description of different generative network models.

| GNM | Description of wiring rule |
| --- | --- |
| *Spatial* | Spatial distance between regions (i.e., wiring cost) |
| *Gene co-expression* | Inter-regional similarity in gene expression combined with a wiring cost |
| *Receptor similarity* | Inter-regional similarity in receptor density combined with a wiring cost |
| *Laminar similarity* | Inter-regional similarity in cortical layer histology combined with a wiring cost |
| *Metabolic coupling* | Inter-regional similarity in PET timeseries combined with a wiring cost |
| *Haemodynamic coupling* | Inter-regional similarity in fMRI timeseries combined with a wiring cost |
| *Electrophysiological coupling* | Inter-regional similarity in MEG timeseries combined with a wiring cost |
| *Temporal similarity* | Inter-regional similarity in fMRI timeseries features combined with a wiring cost |
| *Random similarity* | Inter-regional similarity in randomly generated spatially autocorrelated features combined with a wiring cost |
| *Matching* | Inter-regional similarity in topological neighborhoods combined with a wiring cost |

Synthetic networks generated by each GNM were compared to a consensus 200 region left hemisphere network, constructed using diffusion tractography from 326 unrelated participants of the Human Connectome Project (Hansen et al., 2023). Topological similarity between model and empirical connectomes was assessed via the max(KS) statistic – which is the maximum Kolmogorov-Smirnov (KS) statistic between the node degree, node betweenness, node clustering, and connection length cumulative distribution functions for a pair of networks (**Figure 1C**; see **Methods**). Model parameters were fitted by optimizing the max(KS) statistic. Topographical similarity was measured by connection recovery (*R*), defined as the proportion of empirical connections that the model network successfully recaptured, and the Spearman correlation (*ρ*) between model and empirical network’s nodal degree (**Methods**).

### GNMs capture topological, but not topographical properties of empirical networks

The best fit to empirical network topology, indicated by lowest mean max(KS), was achieved by models based on temporal (M = 0.159, SD = 0.016), laminar (M = 0.172, SD = 0.015), and random node similarity (M = 0.181, SD = 0.01; Figure 2A, B). The *matching* GNM (M = 0.197, SD = 0.06) could produce networks with the lowest max(KS) values (0.105), but not consistently. All models achieved mean max(KS) < 0.4, with values closer to 0 indicating a greater topological fit (Figure 2).

**Figure 2.**
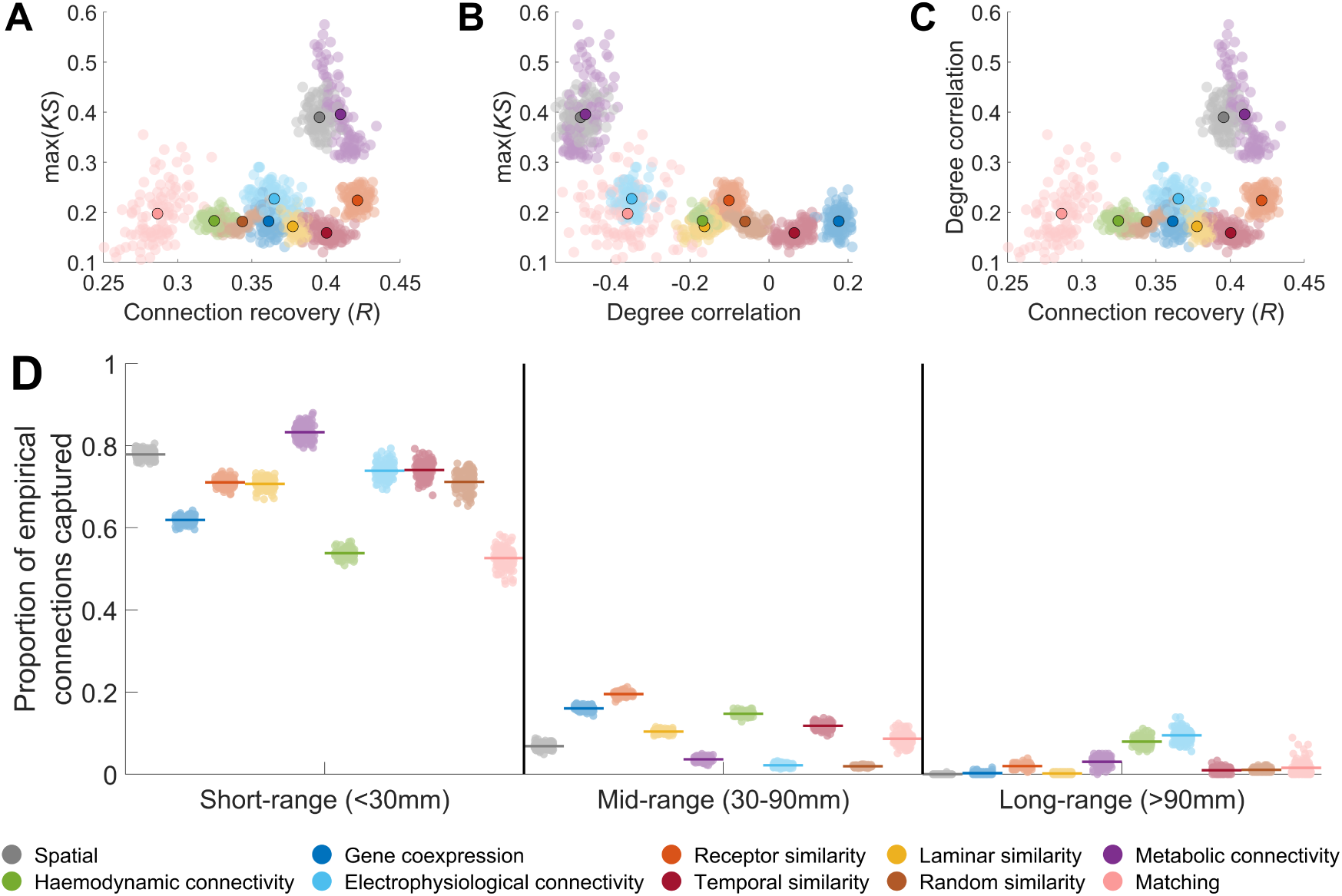
Performance of generative network models in capturing topological and topographical properties. **(A)** Relationship between max(KS) and connection recovery of synthetic model networks. **(B)** Relationship between max(KS) and the correlation between empirical and model node degree. **(C)** Relationship between connection recovery and the correlation between empirical and model degree. In each plot, the outlined point indicates the average across model runs for different model formulations (**D**) Proportion of empirical short-range, mid-range, and long-range connections captured by the best fitting generative network models. The coloured line indicates the average overlap, while each point indicates the result for an individual model network.

The best max(KS) value we observed was in line with previous observations using GNMs (Akarca et al., 2021; Arnatkevičiūtė et al., 2021; Betzel et al., 2016; Oldham, Fulcher, et al., 2022). Despite this comparable performance in capturing network topology, the mean connection overlap between empirical and synthetic networks ranged between only 32% and 42% across all GNM formulations (Figure 2A). Similar performance was observed across alternative GNM formulations (**Figure S1-S2**). Degree topography was also poorly captured, with Spearman correlations between empirical and model degree sequences ranging from – 0.50 to 0.18 (Figure 2B-C**; Figure S1-S2**).

We observed a strong distance-dependence in connection recovery. Short-range empirical connections (<30 mm) were well captured (51–84%), but recovery dropped to 2– 20% for mid-range (30–90 mm) and 0–10% for long-range (>90 mm) connections (Figure 2B**; Figure S3-S4**). Examining the false discovery rate of connections formed by the GNMs (i.e., the proportion of generated connections which were not in the empirical data), further indicated that the majority (>80%) of all connections formed at mid- and long-distance scales were not observed empirically (**Figure S5**). These findings suggest that GNMs can produce synthetic networks with similar topological properties to empirical connections but fail to accurately model long-range connections, leading to poor topographical correspondence between model and data. We also examined the properties of networks produced by the GNMs that had the highest degree sequence correlations. As expected, these networks had a slightly better degree topography (*ρ* ≈ 0.3), but connection recovery remained low (≈25%; **Figure S6**), and long-range connections were still poorly captured (**Figure S7**).

Long-range connections are essential for defining nodal degree topography. To demonstrate this, we applied a progressive connection-length thresholding procedure to the empirical network. When retaining only short-range connections (<30 mm), the degree sequence was only weakly correlated with the unthresholded network (*r* = 0.23). As longer connections were incrementally added, correlation with the original degree sequence increased substantially (Figure 3). In contrast, removing short connections (<60 mm) had minimal impact on degree topography (**Figure S8**), indicating that long-range connectivity— especially among hub regions—plays a central role in shaping cortical degree distribution, despite representing only ≈26% of all connections.

**Figure 3.**
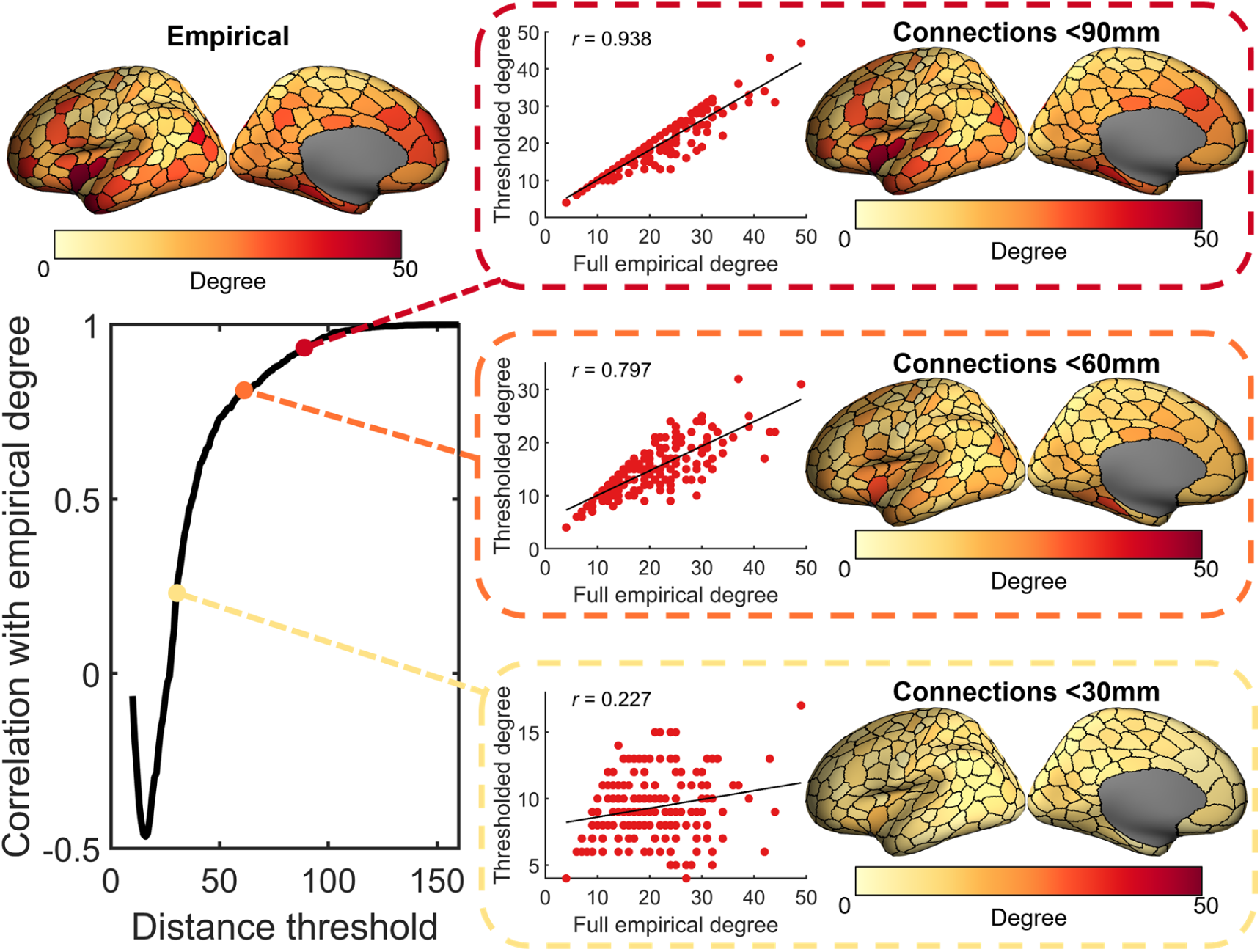
Similarity of the empirical degree sequence at different distance thresholds. Degree is calculated only using connections with a length less than the current distance threshold. The thresholded degree is then correlated (Pearson correlation) with the full empirical degree (i.e., degree calculated when using all empirical connections). Degree is largely shaped by connections >30mm (particularly mid-range connections), meaning that these specific connections need to be captured to obtain the empirical spatial embedding of nodal degree.

Finally, while our primary focus was on left-hemisphere models, we also tested the performance of the 10 primary GNMs on whole-brain networks. Results were broadly consistent with the single-hemisphere findings: max(KS) values ranged from 0.16 to 0.46, and degree correlations from –0.39 to 0.11. However, connection recovery was lower overall (0.16 < *R* < 0.41; **Figure S9**). Some whole-brain GNMs captured more long-range connections than their hemisphere-specific counterparts (e.g., gene co-expression: 31%; receptor similarity: 31%; random similarity: 39%), with this increase being driven by the whole-brain GNMs ability to capture long-range interhemispheric connections between homotopic regions (**Figure S10**). Even so, no GNM could recapture more than 40% of long-range connections.

To determine why GNMs do not accurately model long-range connections between hub nodes, we calculated the probability of each connection (P_ij_) during model fitting as a function of connection length. In all models, P_ij_ was negatively associated with connection length (Figure 4A**; Figure S11**). While at short distances, almost all edges assigned high probability by the model are connected in the empirical networks, at longer ranges only a small proportion of high probability edges are connected (Figure 4B). At connection lengths over 60mm, only 6% of the top 25% most probable connections exist in the empirical data across all GNMs, a similar proportion to that expected by selecting edges at random (Figure 4B). As such, GNMs are highly unlikely to accurately identify the long-range connections between empirical hub nodes (**Figure S12**).

**Figure 4.**
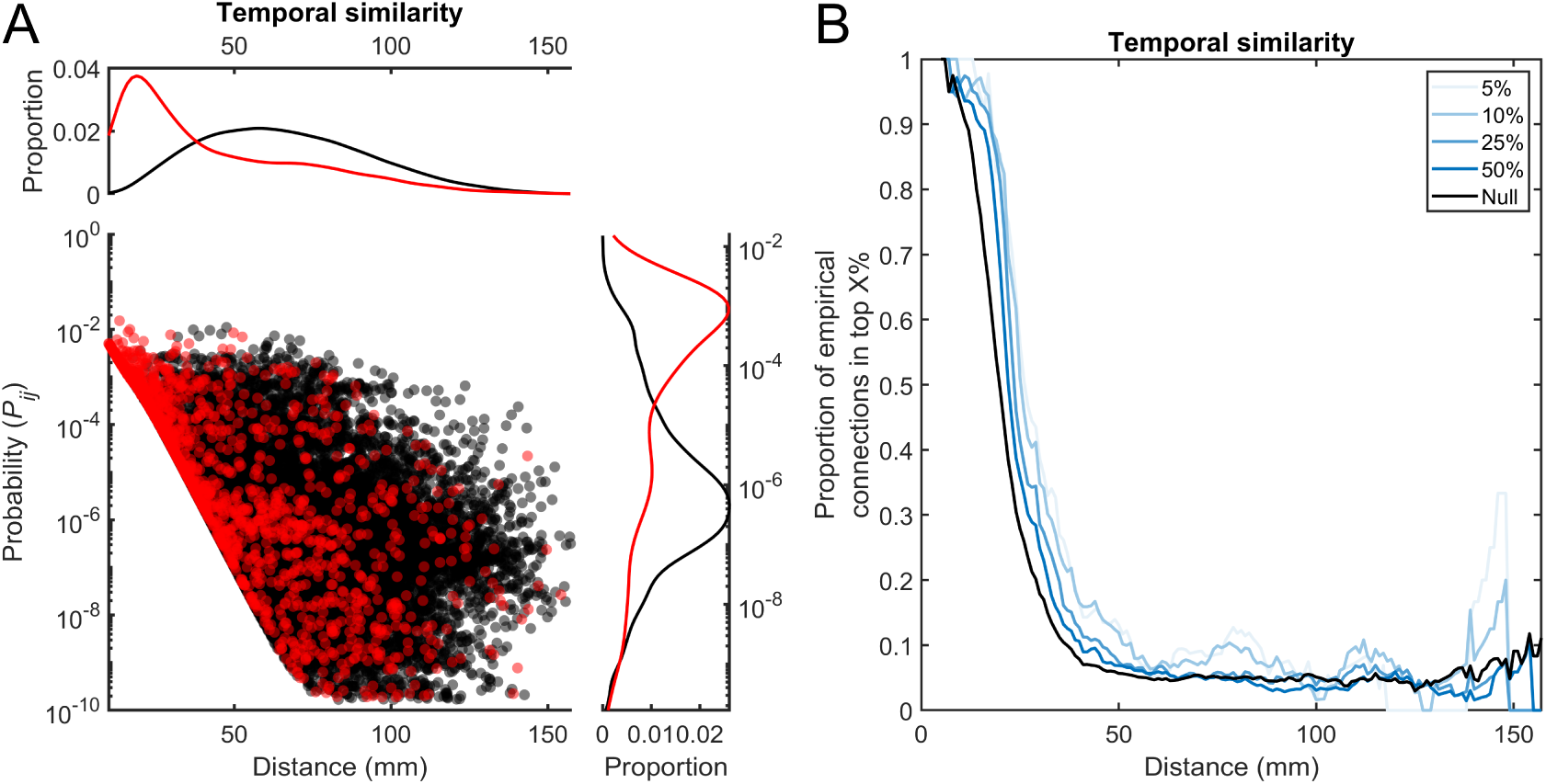
Connection probabilities of the model with the lowest mean max(KS). **(A)** Connection probabilities for the temporal similarity generative network model. The connection probabilities are the mean probability assigned to each connection across all steps of the model. Red points indicate empirical connections (i.e., connections that exist in the empirical data) while black points indicate empirical non-connections (i.e., connections that do not exist in the empirical data). Kernal density plots for the structural and non-structural connections are shown for the distributions of distance/length and probability against the respective axis. **(B)** Most probable connections at different distances. Using a sliding window analysis, at each distance (±5mm) the top X% of connections with the highest probability are found. The proportion of these which are empirical connections is then calculated. The “null” line indicates the proportion of empirical structural connections that exist at that distance (i.e., the probability of selecting an empirical structural connection if all connections at a given distance were equally probable).

### The max(KS) fit statistic can obscure important differences in network structure

Our results have demonstrated that maximising topological similarity between two networks through minimisation of the max(KS) statistic can result in widely different network topographies and poor correspondence between empirical and synthetic network connections (Figure 2A-C). GNMs were unable to recreate the empirical topographic layout of degree or long-range connections, yet the topological similarity of empirical and synthetic connectomes as measured by the max(KS) statistic remains high. To understand this disconnect, we sought to quantify how variations in the max(KS) statistic relates to differences in network topology and topography. To do so, we incrementally rewired connections in the empirical human brain network—either at random or based on connection length—and measured max(KS) between the rewired and original networks (see **Methods**, Figure 5).

**Figure 5.**
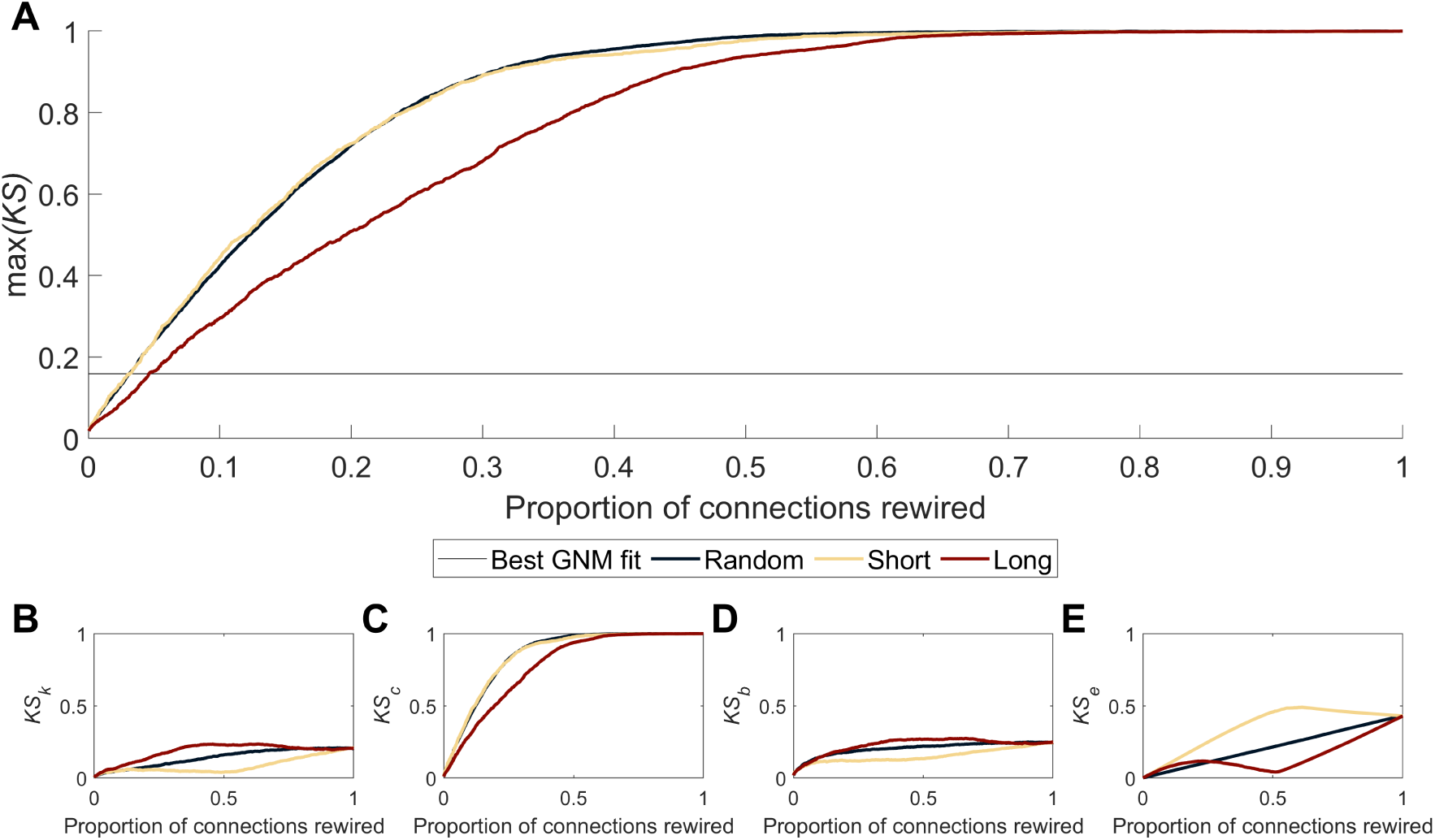
Effect of iteratively rewiring connections on max(KS). **(A)** The proportion of connections rewired and its effect of max(KS) for different rewiring algorithms. The *random* algorithm rewired connections at random but of a similar length; *short* rewired connections from shortest to longest connections with those of a similar length and; *long* rewired connections from longest to shortest connections with those of a similar length. The dotted line indicates the lowest max(KS) fit obtained in the GNMs. Plots also show the change in the KS statistics for the degree **(B)**, clustering **(C)**, betweenness **(D)**, and connection length **(E)** distributions (KS_k_, KS_c_, KS_b_, and KS_e_ respectfully). Changes in max(KS) were almost exclusively driven by changes in clustering.

We observed that max(KS) increased rapidly as connections were rewired (Figure 5A), such that rewiring of just 62 connections resulted in a max(KS) of 0.16 with 97% of empirical connections retained (Figure 5A). This value is comparable to the max(KS) of the best performing GNMs (Figure 2), which only replicated at most 40% of connections.

Constraining the algorithm to only rewire long connections, thus leaving shorter connections in place, reduced this effect with max(KS) most sensitive to changes in shorter connections (Figure 5A). When we examined how the correlation between the nodal degree of the original and rewired network varied across iterations, we found that rewiring long-range connections caused a greater decrease in the correlation than rewiring short-range connections (**Figure S13**).

The max(KS) statistic is a composite of four different topological properties (degree, clustering, betweenness, connection length; Figure 1). Examining the KS statistic for each of these four properties, we found that the increased max(KS) largely reflected changes in the network clustering coefficient (Figure 5B-E). The rewiring analysis indicates max(KS) is highly sensitive to small perturbations in network connectivity that disrupt network clustering.

While the pattern of topological and topographical features is relatively conserved across individual empirical brain networks, there is still individual variation. To establish how this variation is reflected in variations of the max(KS) statistics in empirical connectomes, we compared 972 individual networks constructed from Human Connectome Project data, using 10 parcellations (Schaefer 100 to 1000) and deterministic or probabilistic tractography (see **Methods**). All networks produced using a given parcellation and tractography algorithm were matched for the same density. We fitted the 10 GNM types to the ensemble of individual empirical networks generated with the Schaefer 400 parcellation. For each individual, we a) identified the synthetic network with the lowest max(KS) value and b) calculated max(KS) with respect to every other empirical network as well.

We found that the similarity between empirical brain networks, as measured by the max(KS) statistic, varied depending on parcellation size, with finer parcellations showing greater consistency across individuals (Figure 6). Overall, max(KS) varied from 0.035 to 0.325 between individual empirical network pairs, with connection overlaps ranging from 0.3 to 0.85%.

**Figure 6.**
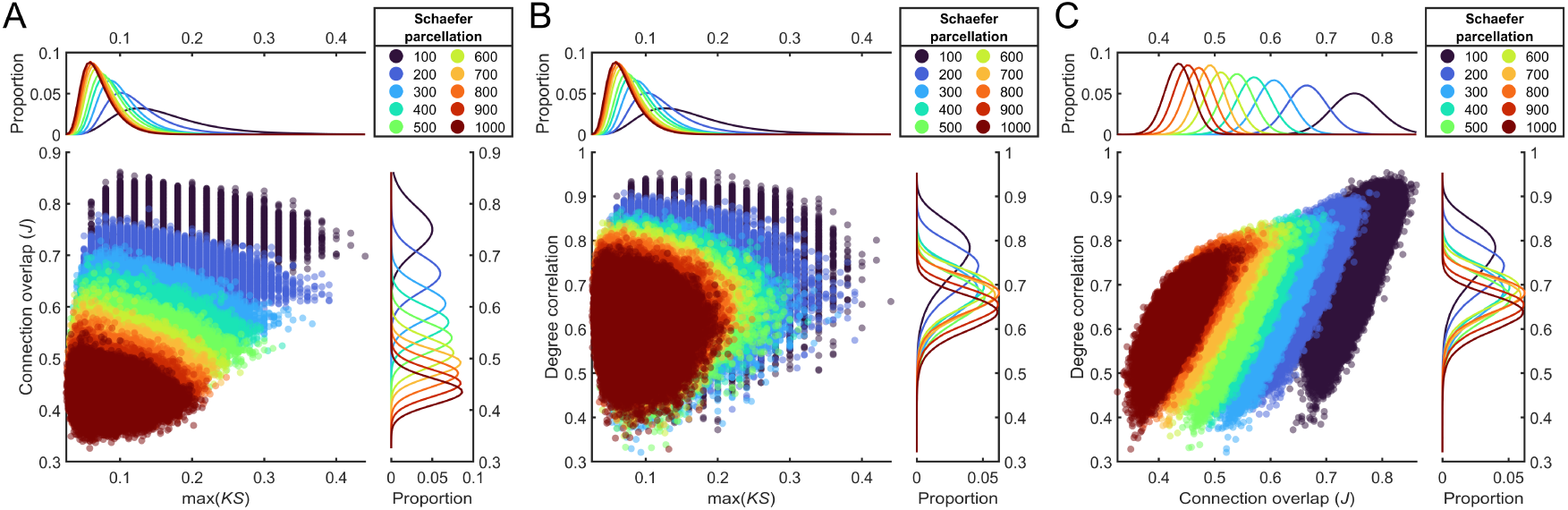
Performance of max(KS), connection overlap, and similarity of the degree distribution when comparing empirical networks produced using probabilistic tractography. **(A)** Relationship between max(KS) and connection overlap. **(B)** Relationship between max(KS) and the correlation between two empirical networks nodal degree. **(C)** Relationship between connection overlap and the correlation between two empirical networks nodal degree. The kernel density plots show the distributions for each parcellation on the respective feature.

Both topological and topographical similarity varied with parcellation resolution. For example, the Schaefer 100 parcellation had a mean max(KS) of 0.15 (SD = 0.039) compared to the Schaefer 400 parcellation (0.09, SD = 0.028) and the Schaefer 1000 parcellation (0.070, SD = 0.022). Empirical networks created using coarser parcellations also showed greater connection overlap between individuals (e.g., Schaefer 100: *M* = 0.752, SD = 0.025; Schaefer 400: *M* = 0.571, SD = 0.024; Schaefer 1000: *M* = 0.435, SD = 0.021), in addition to higher degree correlations (e.g., Schaefer 100: *M* = 0.765, SD = 0.065; Schaefer 400: *M* = 0.685, SD = 0.052; Schaefer 1000: *M* = 0.63, SD = 0.043), than finer resolution parcellations (Figure 6). Similar results were obtained when using deterministic tractography (**Figure S14**).

On average, topological similarity (as measured by max(KS)) between individual empirical and synthetic networks generally occupied a similar range to inter-individual similarity, with over 94% of *matching*, *gene similarity*, and *temporal similarity* GNM networks within the range of interindividual empirical-empirical values (other models showed 6-83% overlap; Figure 7A). However, for measures of topography (i.e., connection overlap and degree correlation), empirical-synthetic network comparisons showed almost no overlap with the range of values recorded in empirical-empirical networks (Figure 7B). Overall, this result demonstrates that topographic similarity can show wide variation even within a constrained range of max(KS) values.

**Figure 7.**
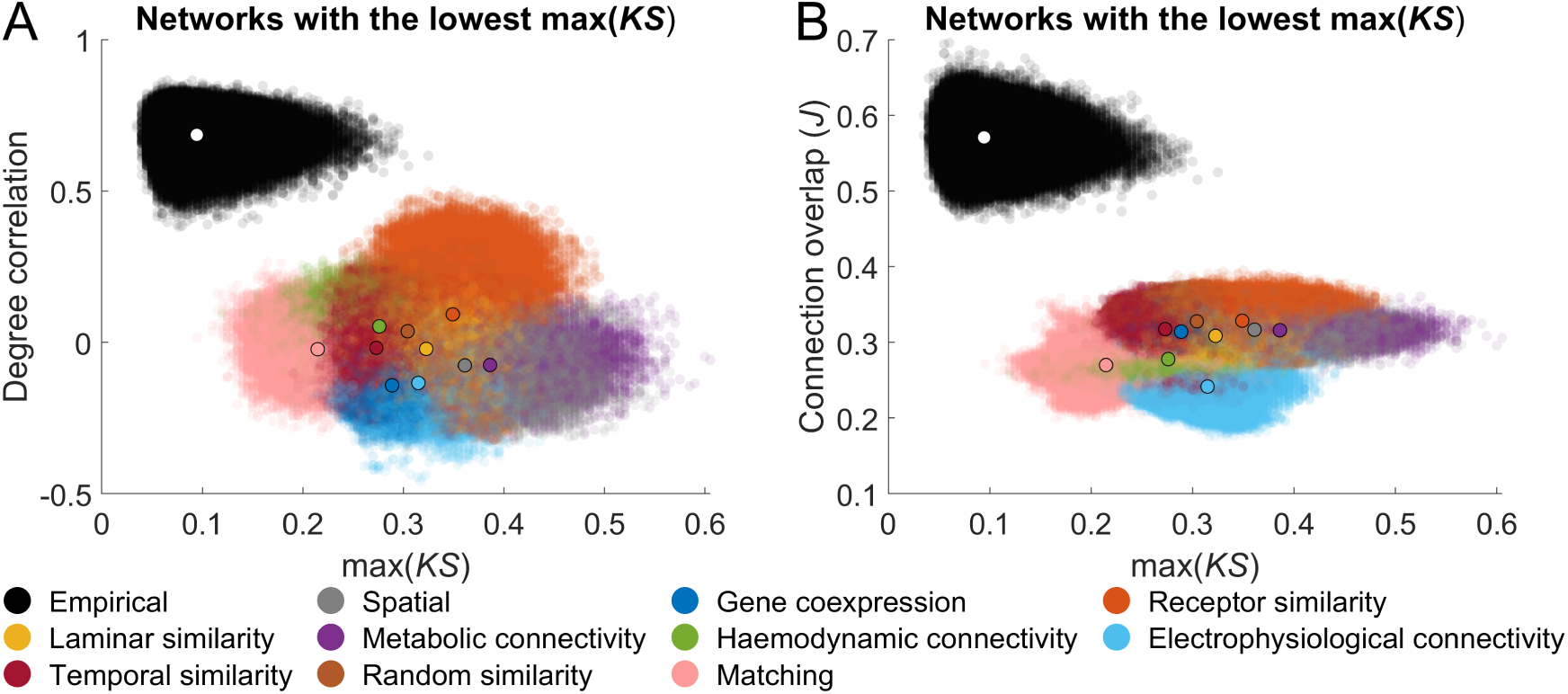
Performance of max(KS) in empirical networks. Scatter plots depict the relationship between max(KS) and the **(A)** degree correlation or **(B)** connection overlap for both empirical-synthetic (GNM-produced vs. empirical networks) and empirical-empirical (empirical network vs. empirical network) comparisons. In each plot, the black-outlined dot represents the mean value for each models empirical-synthetic comparisons, with the white dot indicating the mean for empirical-empirical comparisons. The networks with the lowest max(KS) value for each participant were extracted, and their similarity to all other empirical participant networks was computed. Additionally, all pairs of individual empirical networks were compared using each similarity metric. All empirical networks were generated via probabilistic tractography and using the Schaefer 400 parcellation.

### Different similarity measures do not identify more topographically similar networks

Our analysis reveals that the max(KS) statistic is highly sensitive to small changes in short-range connectivity (Figure 5), but is not sensitive to large differences in long-range connectivity. We next assessed if alternative measures of network similarity could identify networks with greater topological and topographic similarity to empirical data (**Table S2-S3**). These alternatives were informed by approaches proposed by other studies (Akarca et al., 2022; Goñi et al., 2013; Liu et al., 2024), including TND (similarity across global measures of topology), TF_diff_ (Euclidean norm of the difference between correlation matrices of topological features), max(r_d_) (maximum Pearson distance across degree/clustering/betweenness/connection length), and max(RMSE) (maximum mean-root-squared error across degree/clustering/closeness/connection length).

We first compared how the different similarity measures performed in the rewiring analysis, finding that the alternative measures showed a rate of change equal to or lower than that of max(KS) (**Figure S15**). Next, we computed the similarity of each network generated during the optimisation using these alternative measures. While the optimisation procedure was intended to maximise max(KS), as this procedure densely samples the entire parameter space, it allows for exploration if other parameters can produce a network similar to the empirical data. For each individual empirical network a) the most similar network (under each metric) generated during the optimisation was selected, and b) the similarity of this network to all other individual empirical networks on given measure was calculated. As with max(KS), no alternative metric identified synthetic networks with both a high degree correlation and connection overlap to empirical data (Figure 8). Some measures— max(r_d_) and max(RMSE)—indicated that synthetic networks were distinct from empirical networks (Figure 8A, B, E**, F**), (TND, TF_diff_) identified networks that were as similar to empirical networks as empirical networks were to each other (mean overlap across GNMs: TND = 99%; TF_diff_ = 99%; Figure 8C, D, G**, H**). Measures that indicated the generated networks were similar to empirical data primarily assessed topological properties, whereas those that suggested dissimilarity also accounted for topographical differences. None of the alternative measures successfully identified networks with strong topographical similarity to empirical data. Together, these findings suggest that alternative objective functions are unlikely to improve the topographical correspondence between GNM-generated and empirical networks.

**Figure 8.**
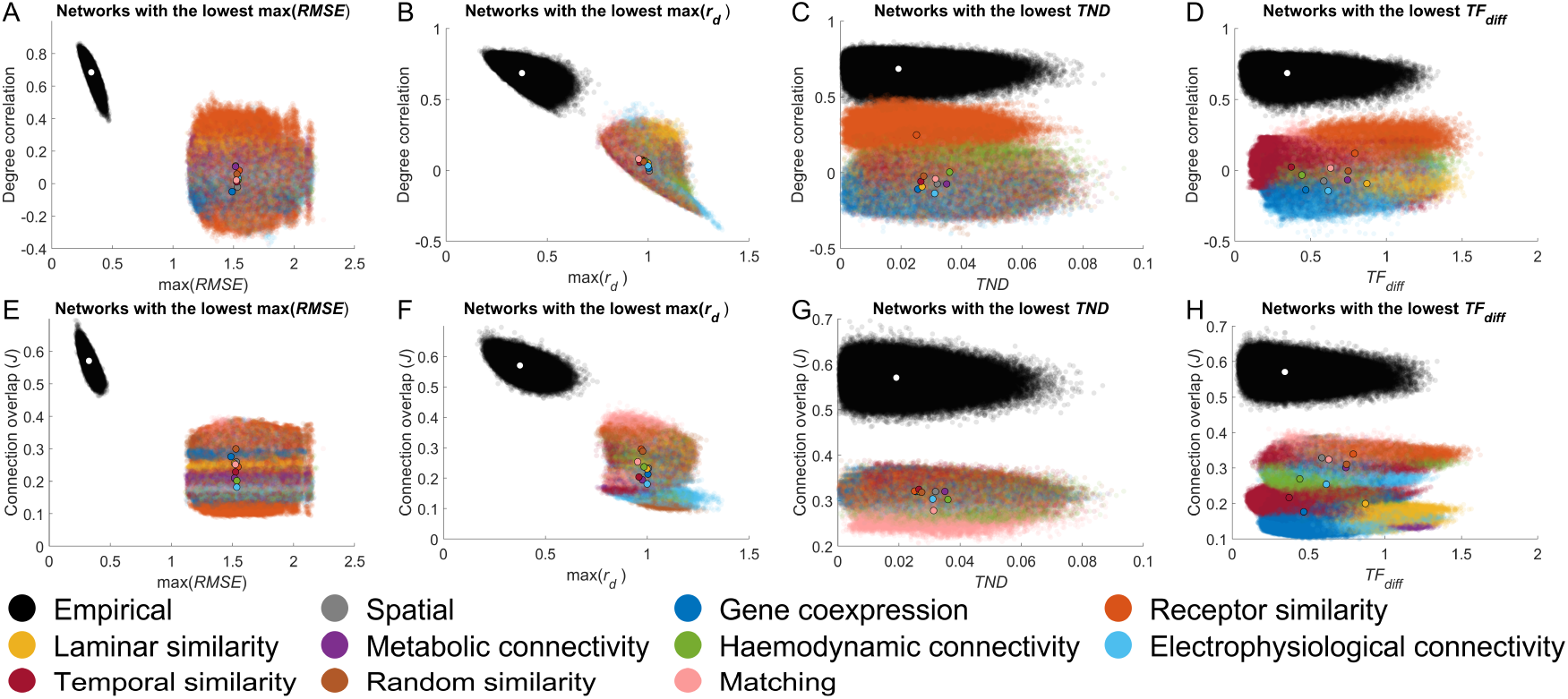
Performance of different similarity measures in generative network models (GNMs) and empirical networks. Scatter plots depict the relationship between degree correlation and **(A)** max(RMSE); **(B)** max(r_d_); **(C)** TND; and **(D)** TF_diff_ for both empirical-synthetic (GNM-produced vs. empirical networks) and empirical-empirical (empirical network vs. empirical network) comparisons. Similarly, scatter plots show the relationship between the connection overlap and **(E)** max(RMSE); **(F)** max(r_d_); **(G)** TND; and **(H)** TF_diff_ for the same comparisons. In each plot, the black-outlined dot represents the mean value for each models empirical-synthetic comparisons, with the white dot indicating the mean for empirical-empirical comparisons. For each similarity measure, the networks with the lowest value for each participant were extracted, and their similarity to all other empirical participant networks was computed. Additionally, all pairs of individual empirical networks were compared using each similarity metric. All empirical networks were generated via probabilistic tractography and using the Schaefer 400 parcellation.

## Discussion

GNMs offer a powerful framework for investigating potential mechanisms and constraints of human brain network organisation. Nonetheless, our results establish that many popular topological GNMs routinely fail to capture critical topographical network organisation due to inaccurate reconstruction of long-range connections between cortical hubs. This is because traditional measures used for model optimization and evaluation based on the similarity of network topology are biased toward the reconstruction of short-range connections. GNMs prioritizing similarity of regional anatomical, cellular, molecular, or dynamical features also fail to overcome this bias, with selection of edges based on similarity in random data showing comparable performance. Our findings therefore indicate that current approaches for defining and fitting GNMs do not identify any single model or parameter combination that can produce a network which accurately captures both the topology and topography of empirical connectome data.

Our analysis indicates that a major reason for the noted failure of extant GNMs to capture connectome topography is their inability to reproduce the specific node-node pairing of long-range connections—a failure that is particularly evident when examining the degree sequence of empirical data (Arnatkevičiūtė et al., 2021; Y. Chen et al., 2017; Oldham, Fulcher, et al., 2022; Zhang et al., 2021), is that the models do not reproduce the specific node-node pairing of long-range connections, which play a critical role in shaping the rich-club organisation of the brain (Arnatkevičiūtė et al., 2021; Fulcher & Fornito, 2016; van den Heuvel et al., 2012; van den Heuvel & Sporns, 2011, 2013). This is because these connections do not have a major impact on the max(KS) statistic, which is the objective function most often used to optimize GNM parameters.

In the GNMs evaluated here, long-range connections impart a significant wiring cost, D_ij_. A high value of inter-regional or topological similarity, F_ij_, is therefore required to overcome this distance penalty and increase the probability of connection. In almost every formulation tested, empirically-connected and unconnected edges could not be clearly distinguished based on a given feature at longer distances (Figure 4A). A partial exception to this was long-range inter-hemispheric connections, who in certain models (e.g., gene co-expression) were highly replicated, likely because homotopic regions have similar biophysiological properties which helps promote long-range connections between them.

However, for most other long-range connections, the extent of biological similarity between unconnected regions was similar to or greater than the similarity of connected regions. This observation appears to stand in contrast to previous studies demonstrating that structurally connected cortical areas are on average more similar to each other than to unconnected areas (Arnatkevičiūtė et al., 2018; Arnatkevičiūtė et al., 2021; Fulcher & Fornito, 2016; Hansen et al., 2023; Hilgetag et al., 2019; Wei et al., 2019), or that hubs regions share similar patterns of gene expression and cortical structure (Arnatkevičiūtė et al., 2021; Paquola et al., 2019, 2020; Sydnor et al., 2021). However the extent of biological assortativity may vary with distance in the brain, with short range connections existing between biologically similar regions and long-range connections being between biologically dissimilar regions (Bazinet et al., 2023; Betzel & Bassett, 2018; Markov et al., 2013). Currently GNMs cannot model such a distance dependency, which may partially account for why long-range connections cannot be accurately captured.

Compounding the issue of GNMs accurately capturing connectivity is that max(KS), the primary measure used to determine network similarity, is sensitive to differences in short-range, but not long-range connectivity. Short-range connections most heavily contribute to defining max(KS), therefore GNMs that can accurately capture the arrangement of short-range connections will perform well on this measure. This accounts for why many models achieve a low max(KS) as the most probable connections are short-range, even when based on randomly spatially autocorrelated features (Figure 4C**; Figure S8**). This also means that different GNMs can produce very different networks to each other, yet be gauged as equally similar to the empirical data merely because they have accurately captured short-range connections. Conversely, if a network accurately recaptures long-range connectivity, and therefore likely topographical properties, but not short-range connectivity it will do poorly on max(KS). In either case, it is clear that max(KS) does not provide a complete picture of the data, thus using it as an objective function on it to identify the most similar network to the modelled data may therefore result in overly optimistic estimates of the extent to which a given model captures desired empirical properties. Even if current GNMs were adapted to use an objective function sensitive to topography (e.g., max(RMSE)), our results suggest they would still fall short of accurately reproducing the full topography of empirical connectomes, due to fundamental limitations in the model framework itself.

Even though for the current class of GNMs an alternative objective function is unlikely to dramatically improve their performance, future work should take care when selecting an optimisation function as this decision is non-trivial (Betzel & Bassett, 2017). For example if the objective function was defined as connection overlap, this could result in a synthetic network that accurately recaptures many empirical connections but has a radically different topology (Betzel et al., 2016; Betzel & Bassett, 2017). However, our results show that the opposite is also true, two networks can have a very similar topology despite few overlapping edges. Objective functions are selected on the basis they are capturing key relevant properties of the network (Betzel et al., 2016; Betzel & Bassett, 2017). The class of GNMs used in this paper were developed to provide a parsimonious description of the connectomes organisation with respect to its topology, therefore the objective function (i.e., max(KS)) was designed to reflect this (Betzel et al., 2016). It is unlikely that optimising for topology will produce a network with a prescribed topography, as a network with a given topology can have a large number of different topographical representations (Kaiser & Hilgetag, 2006). While important, topology alone cannot explain the entire organisation of the connectome. The spatial embedding of connectivity dictates the function of brain networks (Gollo et al., 2018; Kaiser & Hilgetag, 2006; Pang et al., 2023; Roberts et al., 2016), for example specific long-range connections are needed to promote brain-like dynamics (Vohryzek et al., 2025). Therefore, network topography is a crucial organisational property which GNMs should aim to capture. If GNMs are to be used to make stronger claims about the mechanisms and constraints of connectome organisation, the objective function will likely need to be amended to reflect topographical similarity.

### Improving generative network models

Our findings indicate that current GNMs offer an insufficient account of both the topology and topography of human connectomes mapped with diffusion MRI. It is useful to reflect on the potential reasons for this failure as a way of improving the biological accuracy of the models.

One possible factor limiting the accuracy of these models is that they fail to account for regional differences in the timing with which different connections form, a phenomenon known as developmental heterochronicity (Beul et al., 2018; Goulas, Betzel, et al., 2019; Goulas, Majka, et al., 2019; Hilgetag et al., 2019). Differential timing of cortical network development has been suggested as a mechanism underlying the emergence of network features like hubs and long-range connections (Kaiser, 2017; Oldham, Ball, et al., 2022; Oldham & Fornito, 2019). Prior work indicates that incorporating global developmental changes in geometry can improve model fits (Oldham, Fulcher, et al., 2022), but imbuing GNMs with regionally patterned windows within which specific connections are formed may be critical for shaping the degree topography of the human brain. Indeed, it has been noted that areas with poorer laminar differentiation complete neurogenesis and start forming connections earlier than areas with more pronounced differentiation (Beul et al., 2017; Cahalane et al., 2012; Epihova et al., 2024; Finlay & Uchiyama, 2015; Scholtens et al., 2014; van den Heuvel et al., 2015). Since network hubs tend to be located in transmodal regions that show poorer differentiation than primary sensory cortices (Buckner et al., 2009; Margulies et al., 2016), they may obtain an early advantage that drives a rich-get-richer phenomenon known to play a critical role in the emergence of network hubs in a diverse array of systems (Barabási & Albert, 1999; Nicosia et al., 2013).

Furthermore, the best way to parameterize inter-regional variability in biophysiological features remains unclear. Although cytoarchitectonic similarity has been proposed as a key criterion for predicting if two regions are connected (Arnatkevičiūtė et al., 2021; Barbas, 2015; Beul et al., 2017, 2018; Hilgetag et al., 2019; Oldham, Fulcher, et al., 2022), some evidence suggests that dissimilar regions may be more likely to be connected across long distances (Bazinet et al., 2023; Betzel & Bassett, 2018; Markov et al., 2013), models may need to account for a distance-dependent homophilic attachment mechanism. In some GNMs based on biophysiological similarity, inter-hemispheric connections could be recaptured with greater accuracy than long-range intra-hemispheric ones, suggesting that long-range inter- and intra-hemispheric connectivity may be governed by different attachment mechanisms. Additionally, our models did not directly examine how homophilic attachment based on biophysiological properties may interact with homophilic attachment based on topological properties. Configuring the models in such a way may expand their ability to capture complex connectivity patterns.

An alternative way forward for GNMs is to move away from models focused on pairwise interactions between regions to capture dynamic properties such as axonal growth (Goulas, Betzel, et al., 2019; Liu et al., 2024; Song et al., 2014). Such models have been implemented for simple 2D or 3D representations of the cortical surface (Goulas, Betzel, et al., 2019; Liu et al., 2024; Song et al., 2014) but could be extended to incorporate the complex geometry of the human brain. These more biophysical models would also have the advantage of easily allowing for connection weights to be modelled, and potentially allow for the shape of the formed tracts to be examined (Liu et al., 2024), providing additional angles with which to evaluate the synthetically produced connectivity.

### Limitations

Several biases have been identified that can limit the biological veracity of empirical connectome data derived from diffusion MRI (Sotiropoulos & Zalesky, 2019). This includes an inherent difficulty in capturing long-range connections (Sotiropoulos & Zalesky, 2019), thus the empirical networks employed in this study may underestimate the true extent of long-range connections in human brain networks. Despite this, GNMs in their current form are still unable to reach this lower bound. Multiple approaches have been developed to mitigate some of the biases in diffusion MRI tractography and network construction (Baum et al., 2018; Oldham et al., 2020), yet variations between methods can still yield networks with major differences in topology and topography (Gajwani et al., 2023; Oldham et al., 2020). While it is possible that observations on GNMs applied to data processed using one pipeline may not generalize to data processed in other ways, we believe that the identified issues with GNMs persist across different processing choices. Prior work using GNMs for networks constructed using deterministic tractography revealed similar findings (Oldham, Fulcher, et al., 2022). We also found that, for networks created using distance-dependent thresholding, which explicitly aims to preserve long-range connections, or strength thresholding, which can decrease estimates of long-range connectivity, GNMs were unable to model the correct positioning of hubs. Furthermore, when we extended our analysis to whole-brain connectivity (as opposed to a single hemisphere) which includes substantially more long-range connections, similar weaknesses were observed. Finally, GNMs have been used to model primate structural brain networks based on tract tracing data (Y. Chen et al., 2017), with similar observations to those obtained in tractography-based human brain networks, most notably a failure to capture hub location and long-range connections.

We did not consider the weighted topology/topography of empirical connectomes, which is a salient feature of brain networks (Fornito et al., 2016; Markov et al., 2011). One recent GNM has been proposed to model connection weights, and shown promise in capturing binary and weighted features (Akarca et al., 2023), although it remains to be seen if the reported model fits, which rely on a similar approach to that investigated here, are subject to the same biases. Our models were restricted to just cortical connections, and thus did not account for the complex connectivity patterns linking the cortex, subcortex, and cerebellum (Müller et al., 2020; Oldham et al., 2025; Oldham & Ball, 2023; Park et al., 2024; Raut et al., 2020; Tian et al., 2020). Expanding GNMs to include all types of brain regions is needed to achieve a more comprehensive understanding of mechanisms underpinning all connectome organisation.

## Conclusion

GNMs can successfully replicate basic topological properties of brain networks but they fall short in capturing topographical properties, particularly the crucial long-range connections observed in empirical data. Additionally, the common practice of evaluating model fits purely based on topology may overlook the true extent of differences between networks. Our findings establish where and why generative network models struggle in capturing brain connectivity and indicate directions for future research to address these challenges and improve model accuracy.

## Methods

### Empirical brain networks

Two empirical datasets were used in this study, a group averaged connectivity matrix (Hansen et al., 2023) and a set of individualised connectivity matrices (Arnatkevičiūtė et al., 2021; Oldham, Fulcher, et al., 2022). Both datasets were constructed from Human Connectome Project (HCP) data (Glasser et al., 2013). The group averaged data had been constructed from the S900 release of the HCP data from 326 participants. In brief, the preprocessed diffusion data had been processed with MRtrix3 using multi-shell multi-tissue constrained spherical deconvolution and tractography was conducted with second-order Integration over Fiber Orientation Distributions (iFOD2). Streamlines were weighted using SIFT2 (Smith et al., 2015), and the group consensus connectivity matrix was created using a distance dependant thresholding algorithm. Further details are available in Hasnen et al., (2023). The group connectivity matrix was binarized for use in this study. The dataset was originally created for the full Schaefer 400, 7 network parcellation (Schaefer et al., 2018), which includes 200 regions for each hemisphere. We primarily used data for the left hemisphere i.e., 200 cortical regions in total as to: a) reduce computational complexity (more nodes/connections in the network increases the time needed to run the GNM) b) reduce any bias that may result of measuring the Euclidean distance from nodes across hemispheres, and c) keep consistent with previous methodologies (Arnatkevičiūtė et al., 2021; Betzel et al., 2016; Oldham, Fulcher, et al., 2022). However, we also examined how a limited number of GNMs performed in recapturing connections/connectivity patterns in the original whole brain network.

The individualised connectivity data had also previously been processed (Arnatkevičiūtė et al., 2021; Oldham, Fulcher, et al., 2022). These data were constructed from the S1200 release of the HCP for 973 participants. A similar series of processing steps were applied as with the first dataset. In addition to iFOD2, a probabilistic tractography algorithm, the deterministic algorithm Fiber Assignment by Continuous Tractography was also run. Further details of the processing steps for the individualised data are available elsewhere (Arnatkevičiūtė et al., 2021; Oldham, Fulcher, et al., 2022). We generated a network for each of the Schaefer 7 network 100-1000 parcellations, using the probabilistic and deterministic tractograms resulting in 20 networks per participant (two different tractograms across 10 different parcellations). As with the group averaged data, we only retained regions in the left hemisphere. One of the 973 individual networks showed noticeably altered connectivity compared to others (i.e., lower density, weaker correlations with other individuals), thus they were excluded from further analysis.

Networks were either unthresholded or thresholded based on connection strength. Thresholding was applied such that only the strongest *E* connections were retained, where *E* was 70% of the minimum number of connection observed for any individual in a given tractography/parcellation combination. While a strength threshold is likely to result in weak, long-range connections being pruned (Fornito et al., 2016), we used this approach to ensure that all individuals for a given network type had the same density with some variation in their topology. Other thresholding approaches involve the creation of a consensus matrix which is then used as a mask for individuals (Betzel et al., 2019; de Reus & van den Heuvel, 2013; Roberts et al., 2017), but such approaches would minimise inter-individual variability in network binary network topology and thus meaning there would be little value in comparing individual binary network organisation.

### Cortical inter-regional similarity measures

We used seven different measures of inter-regional similarity which represent multiple facets of brain organisation (Hansen et al., 2023): (1) gene co-expression (correlation across 8687 genes); (2) receptor similarity (correlation across 18 PET receptor density profiles); (3) laminar similarity (correlation across cortical layer histology); (4) metabolic coupling (correlation in PET timeseries); (5) haemodynamic coupling (correlation in fMRI timeseries); (6) electrophysiological coupling (correlation in MEG timeseries); and (7) temporal similarity (correlation across 6441 fMRI timeseries features). These measures had previously been generated for the Schaefer 400 parcellation and made openly available as part of another study (see Hansen et al., 2023 for details). All measures of inter-regional similarity were provided as a matrix of Pearson correlations. For use with GNMs, each correlation matrix was converted to a Pearson distance (as GNMs cannot tolerate negative values for inter-regional similarity) and then rescaled to the unit interval.

These regional measures are representative of major organisational patterns in the cortex (Hansen et al., 2023; Markello et al., 2022). However, these measures can demonstrate spatial autocorrelations which can explain many properties of the brain (Burt et al., 2020; Markello & Misic, 2021; Shinn et al., 2023). To assess the impact of spatial autocorrelations in GNMs, we created random spatially autocorrelated features and computed inter-regional similarity on those using the following procedure:

1. We generated a random value in the range 0-1 for each cortical region.
2. For each cortical region, we took the average value of each of the nodes it was adjacent to on the cortical surface.
3. We then updated the value of each cortical region to this averaged value (note this was only done once all the averaged values for all regions had been calculated).
4. Steps 2-3 were repeated five times.
5. Steps 1-4 were repeated 20 times to create 20 random spatially autocorrelated features.
6. Random inter-regional similarity was calculated by performing a Pearson correlation across each of these random features for every pair of brain regions.

The above steps were performed for regions of the left hemisphere only. To map the spatially autocorrelated patterns to the right hemisphere (as the parcellation is not homotopic across hemispheres), the left hemisphere random features were mapped to the right hemisphere, and the mean value was taken for each right hemisphere region.

While there are more sophisticated approaches of creating spatially autocorrelated cortical features (Burt et al., 2020), this approach is a fast and efficient way of creating multiple such maps. As with the other measures of inter-regional similarity, the Pearson correlation was converted to a Pearson distance and then rescaled to the unit interval. In addition to these measures of inter-regional similarity, we also examined 12 topologically based models (i.e., topological coupling was between a pair of regions was used for the value of F_ij_; **Table S1**).

Additionally Euclidean distances between all pairs of nodes were calculated to get an estimate of wiring-costs for the GNM. Specifically, the Euclidean distance was taken as the average distance from all the vertices on the cortical surface that were assigned to one region, to all the vertices assigned to another region. Euclidean distances do not account for the curved paths tracts have to take i.e., fibre distances, and so can underestimate the true wiring costs. While more elaborate procedures exist for approximating the fibre lengths and therefore wiring-costs (Oldham, Fulcher, et al., 2022), Euclidean and fibre distances are often strongly correlated (Betzel et al., 2016), and using either to predict connections in brain networks has resulted in similar performance (Rubinov et al., 2015). We therefore deemed Euclidean distances as an acceptable approximation of wiring-costs for within hemisphere connectivity.

### Generative network model formulation

The generative network model adds connections in a sequential manner up to the desired number (i.e., the number of connections in the network the model is attempting to replicate) according to the wiring rule specified in Equation 1. As indicated by this equation, GNMs were primarily formulated as an additive rule, and the distance decay was defined according to an exponential. This additive rule has previously been shown to more accurately capture the trade-offs between model parameters implied by the model than multiplicative formulations (Oldham, Fulcher, et al., 2022). Alternatively, the rule could be formulated as a multiplication of terms

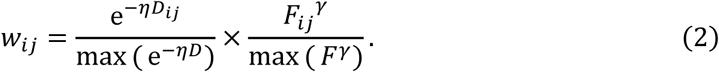

While we primarily model distance as an exponential decay e^-r,Dij^ due to evidence showing the likelihood of a connection existing decreases exponential as a function of its length, other studies have used a power-law decay D_ij_^-r,^ (Betzel et al., 2016), but the specific form of the distance penalty does not change the basic conclusions of our findings.

At each timestep of the model, the weight w_ij_ is converted to a probability according to

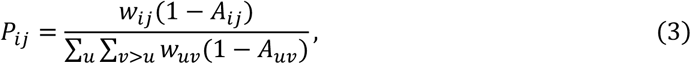

where (1 − A_ij_) ensures that only connections that are not yet present in the network influence the connection probability (i.e., the probability of a connection forming should only be relative to other connections that have not formed yet). At each timestep a single connection is established, with P_ij_ indicating the probability that specific connection will be selected. If F_ij_ is a measure of topology, that measure will be recalculated based on the network will the newly added connection.

As w_ij_ will change at each timestep of the model, so will P_ij_. Therefore, to calculate the overall probability a given connection will form, we average connection probabilities across model timesteps (note the average was only calculated for iterations where the connection had not been added into the network).

### Selecting optimised generative network models

To determine the optimal values for the parameters 1, y, and a for each GNM, we employed a previously developed optimization method (Betzel et al., 2016; Oldham, Fulcher, et al., 2022). The method was implemented as follows:

1. We randomly sampled 2000 points within the parameter space defined by 1, y, and potentially a (if the additive form was used). When an exponential was used, 1 varied between 0 and 2, while for a power-law 1 varied between 0 and 10. For GNMs using inter-regional similarity, y varied between −20 and 200. For those based on topology, y varied between −10 and 10. Additionally, a ranged from 0 to 10 (when α was included as a parameter in the model). These parameter ranges were selected based on initial testing and previous findings (Oldham, Fulcher, et al., 2022).
2. At each sampled point representing a specific combination of 1, y, and a values, we generated a network using the newly defined parameters, resulting in 2000 synthetic networks. We calculated the max(KS) statistic for each generated network. The max(KS) statistic is the maximum KS value (i.e., greatest discrepancy) between the generated and empirical networks in terms of their degree, clustering, betweenness centrality, and connection length distributions.
3. After evaluating all networks, we employed Voronoi tessellation to identify regions (cells) of the parameter space associated with low fit statistics. An additional 2000 points in the parameter space were preferentially sampled from each cell based on the relative probability VC^-/3^, where VC_c_ is the max(KS) of cell c, and /3 controls the likelihood of sampling from cells with low max(KS) values.
4. Steps 2 and 3 were repeated four times, resulting in a total of 10,000 points being evaluated. In each repetition, the probability of sampling cells with better fits was increased by varying /3 from 0.5 to 2 in 0.5 increments, thus converging to an approximate optimum. When the optimisation was applied to the ensemble of individual networks, the network generated by the model for a given parameter combination was compared to all individual networks (Liu et al., 2023). The max(KS) for that parameter combination was taken as the mean of all individual max(KS) values. The networks generated during the optimisation that produced the lowest max(KS) for an individual participant were used for subsequent analyses. When evaluating other similarity measures, we used the networks generated when optimising for max(KS) because (1) there was a greater computational cost to running the GNM again with a different fit statistics and (2), the parameter space is densely sampled so examining from the networks selected is likely to allow the identification of a minimum.

An advantage of this approach is that it allows for the entire parameter space to be adequately sampled and facilitate the identification of a global (approximate) optimum. For the group-averaged consensus network, when the optimal parameters had been identified for a given GNM they were used to generate 100 further networks.

### Rewiring model

Networks were rewired at random, or from the shortest-to-longest connections, or longest-to-shortest. Existing connections could only be rewired/swapped with a non-exiting connection. Once a connection had been rewired, it could not be rewired again. Connections could be rewired with a selected non-existing connection at random, a connection of a similar length, or a connection of dissimilar length. The rewiring continued until all existing connections had been rewired. We ran each rewiring model 30 times and then took the mean max(KS) across these runs.

### Measures of network topology/topography

Details of the various topological/topographical measures used in this study can be found in Tables S1, S2, and S3.

## Code and data availability

All analysis was performed in MATLAB 2023b. Code and data are available from https://github.com/StuartJO/ComingUpShort

## Acknowledgements

Data were provided by the Human Connectome Project, WU-Minn Consortium (Principal Investigators: David Van Essen and Kamil Ugurbil; 1U54MH091657) funded by the 16 NIH Institutes and Centers that support the NIH Blueprint for Neuroscience Research and by the McDonnell Center for Systems Neuroscience at Washington University. S.O is supported by the Brain and Behavior Research Foundation (ID: 31471). A.F. was supported by the Sylvia and Charles Viertel Foundation, National Health and Medical Research Council (IDs: 1197431 and 1146292), and Australian Research Council (IDs: DP200103509). G.B. was support by the National Health and Medical Research Council (ID: 1194497). This research was supported by the Murdoch Children’s Research Institute, the Royal Children’s Hospital, Department of Paediatrics, The University of Melbourne and the Victorian Government’s Operational Infrastructure Support Program. The project was generously supported by The Royal Children’s Hospital Foundation devoted to raising funds for research at The Royal Children’s Hospital.

## Supplementary material for

**Figure S1.**
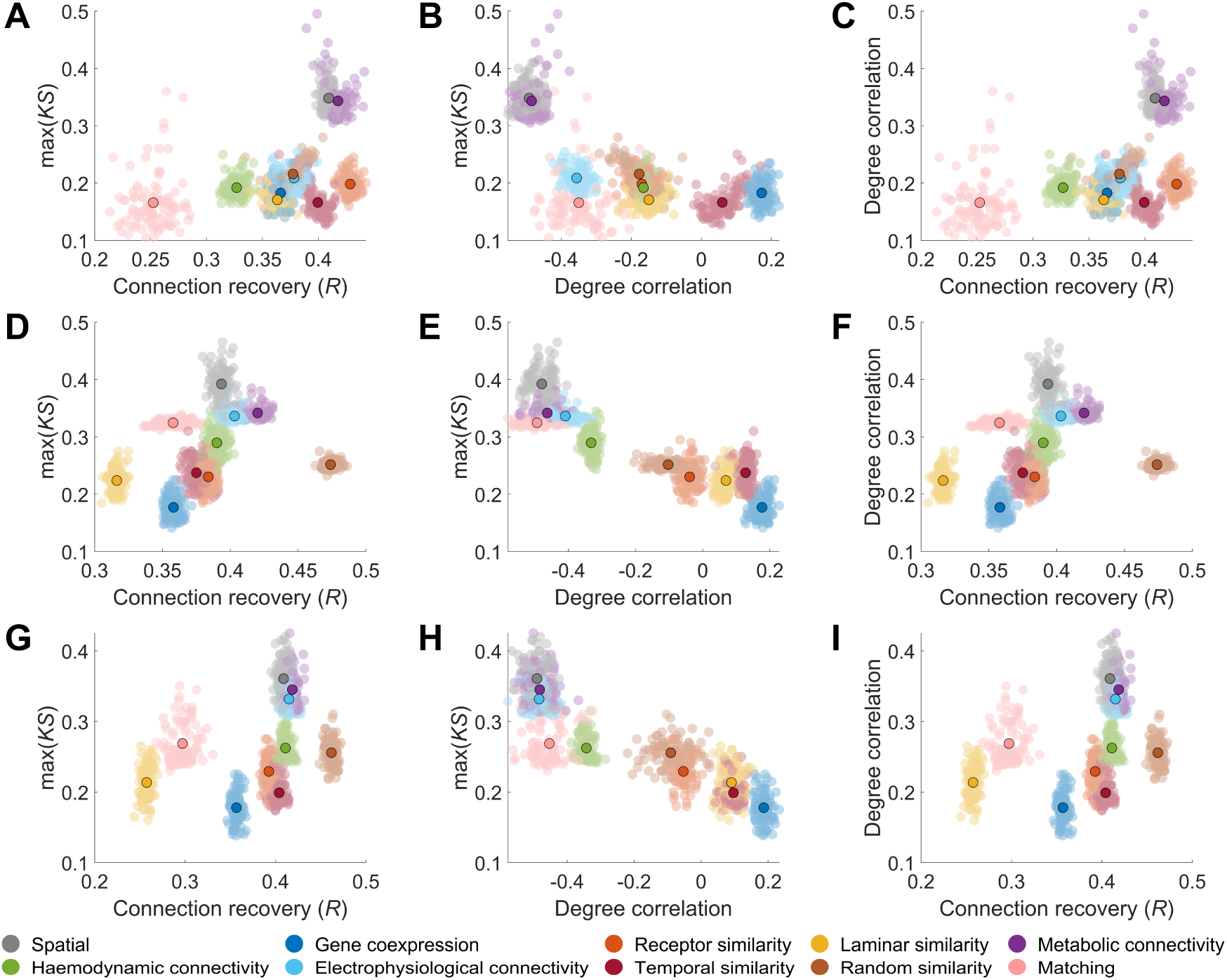
Performance of different formulations of generative network models in max(KS), connections recovered, and similarity of the degree distribution. Generative network models formulated using an additive form and power-law distance decay showing the: **(A)** relationship between max(KS) and connections recovered; **(B)** relationship between max(KS) and the correlation between empirical and model degree; and **(C)** relationship between max(KS) and the correlation between empirical and model degree. Generative network models formulated using a multiplicative form and exponential distance decay showing the: **(D)** relationship between max(KS) and connections recovered; **(E)** relationship between max(KS) and the correlation between empirical and model degree; and **(F)** relationship between max(KS) and the correlation between empirical and model degree. Generative network models formulated using a multiplicative form and power-law distance decay showing the: **(G)** relationship between max(KS) and connections recovered; **(H)** relationship between max(KS) and the correlation between empirical and model degree; and **(I)** relationship between max(KS) and the correlation between empirical and model degree.

**Figure S2.**
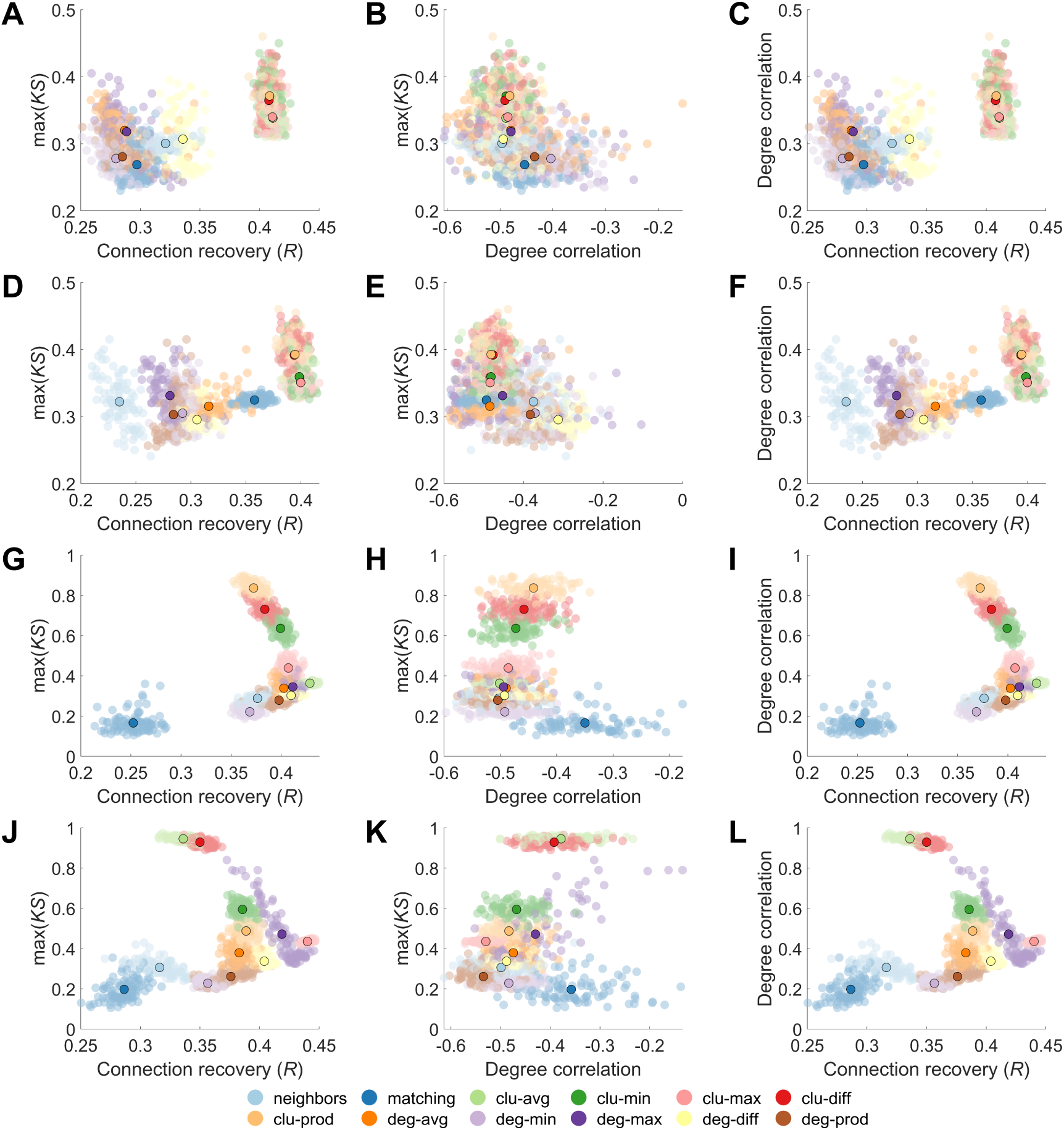
Performance of different topological generative network models in max(KS), connections recovered, and similarity of the degree distribution. Generative network models formulated using a multiplicative form and power-law distance decay showing the: **(A)** relationship between max(KS) and connections recovered; **(B)** relationship between max(KS) and the correlation between empirical and model degree; and **(C)** relationship between max(KS) and the correlation between empirical and model degree. Generative network models formulated using a multiplicative form and exponential distance decay showing the: **(D)** Relationship between max(KS) and connections recovered; **(E)** relationship between max(KS) and the correlation between empirical and model degree; and **(F)** relationship between max(KS) and the correlation between empirical and model degree. Generative network models formulated using an additive form and power-law distance decay showing the: **(G)** relationship between max(KS) and connections recovered; **(H)** relationship between max(KS) and the correlation between empirical and model degree; and **(I)** relationship between max(KS) and the correlation between empirical and model degree. Generative network models formulated using an additive form and exponential distance decay showing the: **(J)** Relationship between max(KS) and connections recovered; **(K)** relationship between max(KS) and the correlation between empirical and model degree; and **(L)** relationship between max(KS) and the correlation between empirical and model degree.

**Figure S3.**
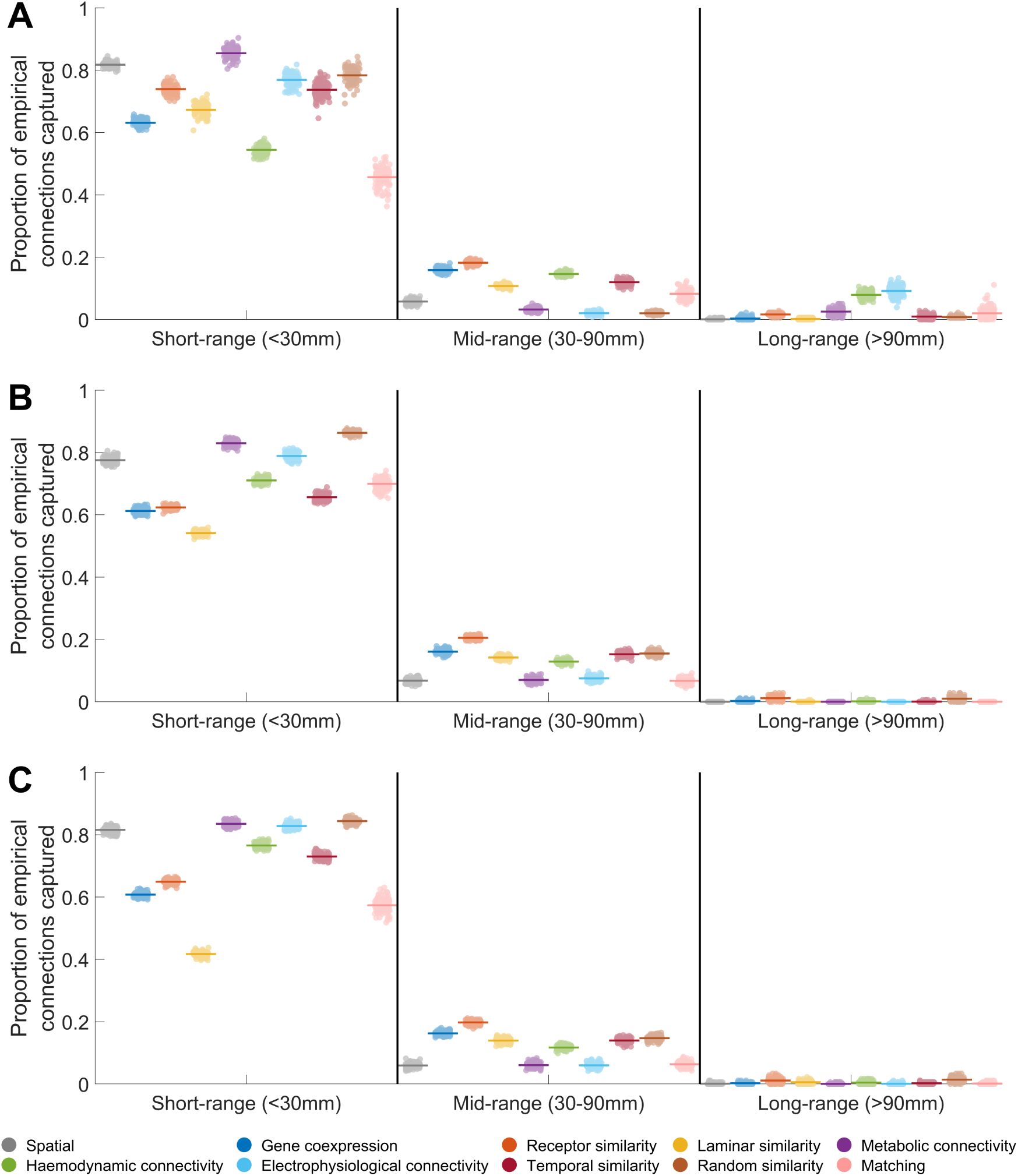
Connections captured at different distance thresholds for different generative network model formulations. Proportion of empirical short-range, mid-range, and long-range connections captured by the best fitting (lowest max(KS)) generative network models for the: (A) additive and power-law distance decay formulation; (B) multiplicative and exponential distance decay formulation; (C) multiplicative and power-law distance decay formulation. The coloured line indicates the average overlap, while each point indicates the result for an individual model network.

**Figure S4.**
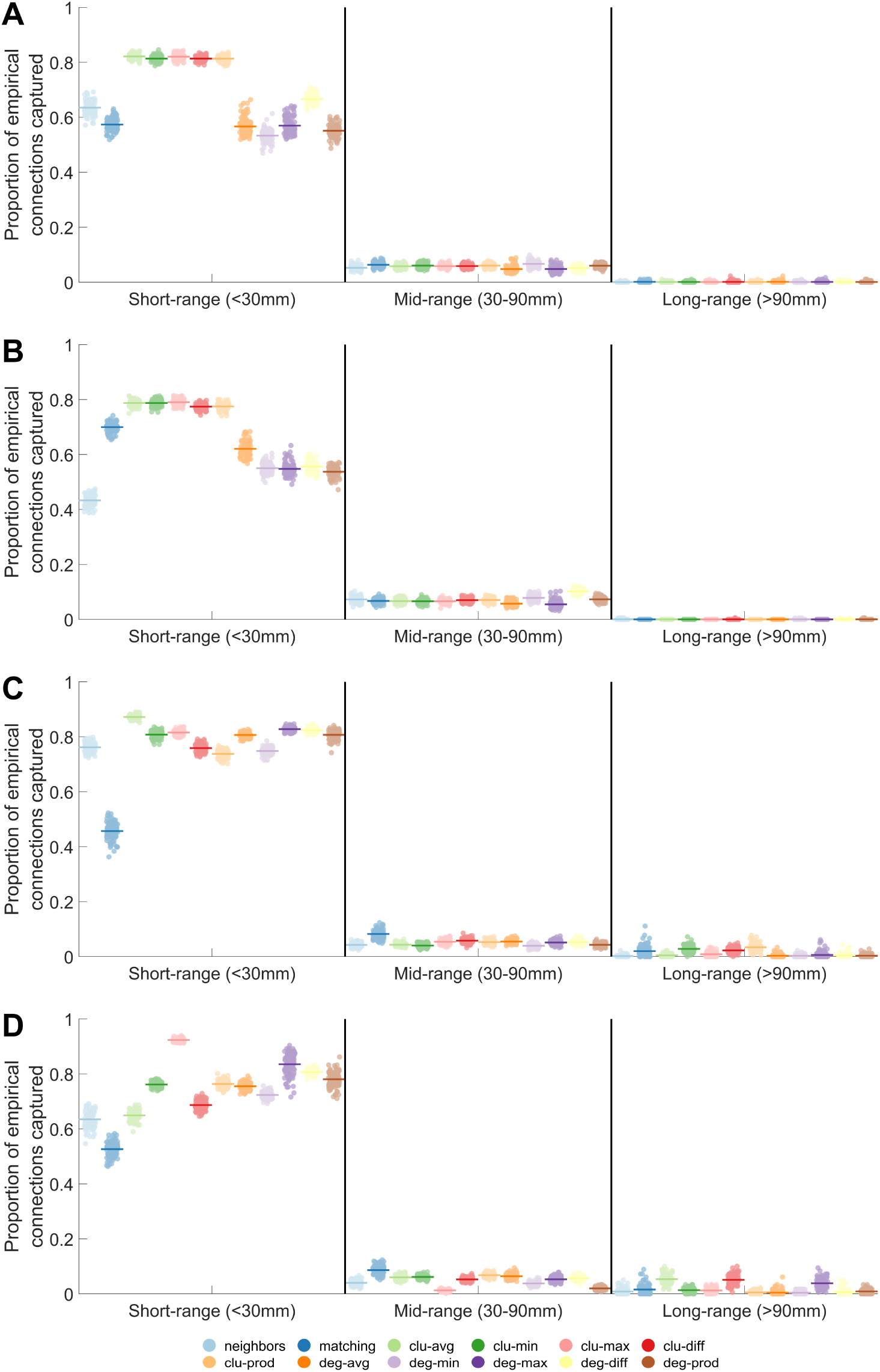
Connections captured at different distance thresholds for topological generative network models. Proportion of empirical short-range, mid-range, and long-range connections captured by the best fitting (lowest max(KS)) topological generative network models for the: (A) multiplicative and power-law distance decay formulation; (B) multiplicative and exponential distance decay formulation; (C) additive and power-law distance decay formulation; (D) additive and exponential distance decay formulation. The coloured line indicates the average overlap, while each point indicates the result for an individual model network.

**Figure S5.**
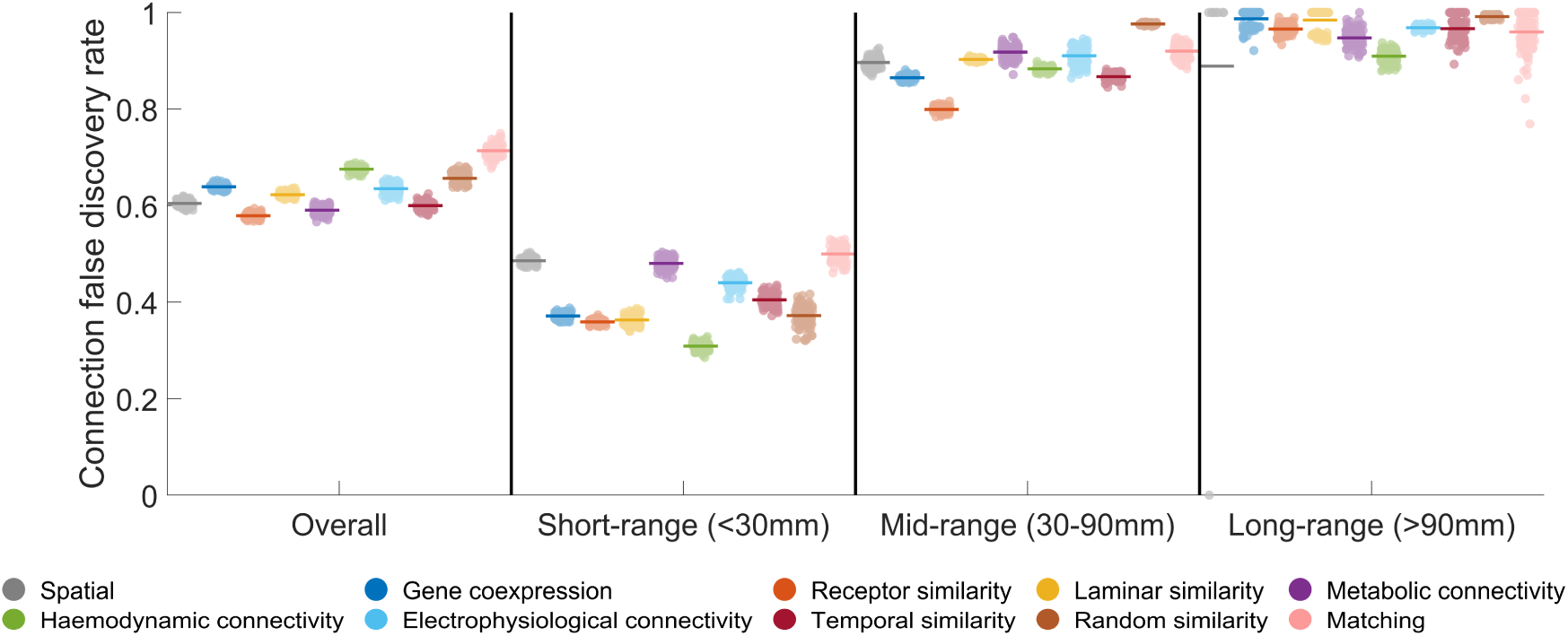
False discovery rate for connections at different distance thresholds. The false discovery rate (i.e., the proportion of connections generated by the model which were not found in the empirical data) is computed for all, short-range, mid-range, and long-range connections for the 10 main GNMs that used the additive, exponential decay formulation. Note that in a single instance of the spatial model, it only generated a single long-range and because that long-range connection existed empirically it achieved a false discovery rate of 0.

**Figure S6.**
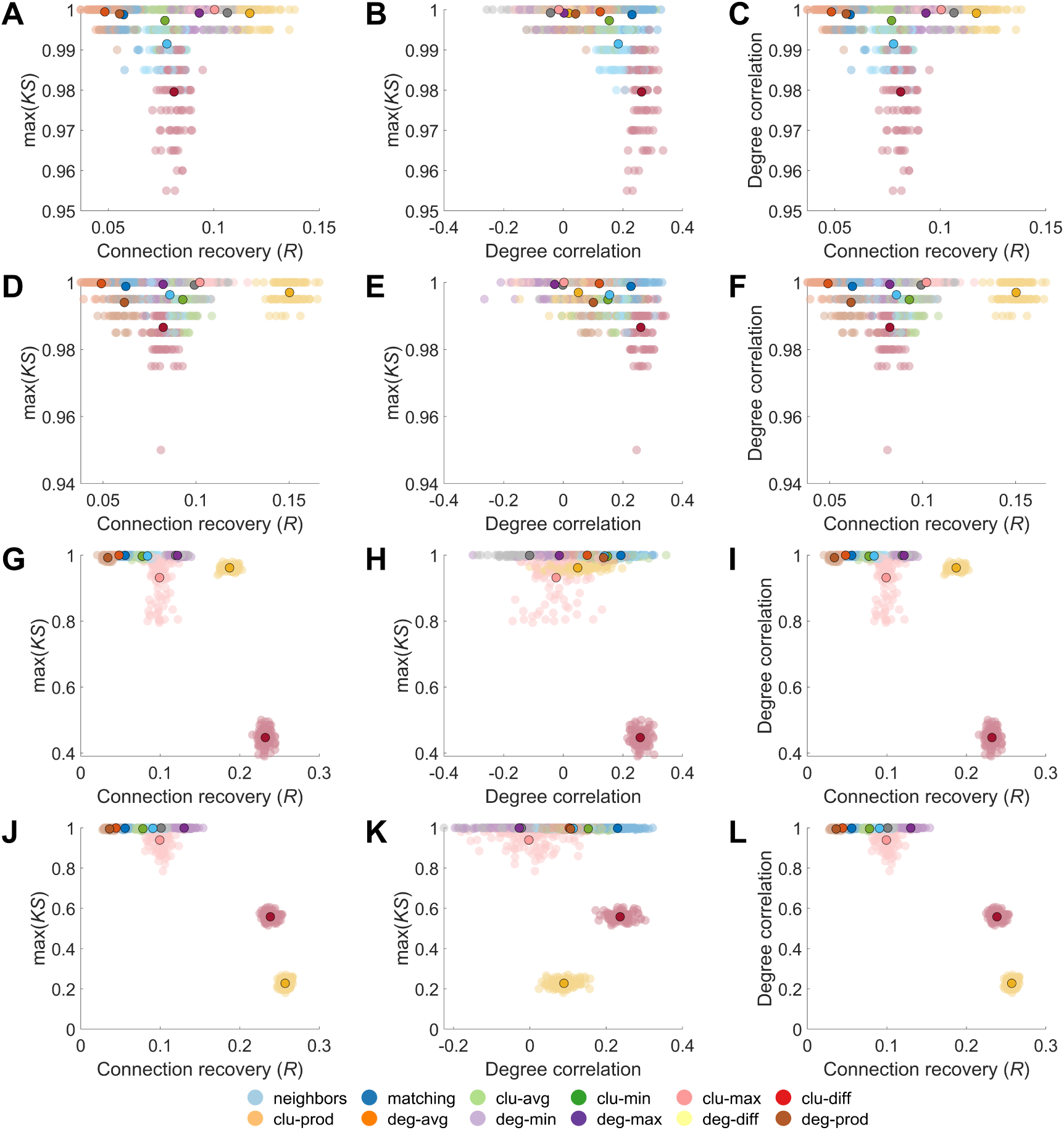
Analysis of generative network models with the strongest degree correlations, evaluated through max(KS), connections recovered, and degree correlation. (**A-C**) Models using an additive form with exponential distance decay: (**A**) Relationship between max(KS) and connections recovered. (**B**) Relationship between max(KS) and the correlation between empirical and model degree. (**C**) Relationship between max(KS) and the correlation between empirical and model degree. (**D-F**) Models using an form with power-law distance decay: (**D**) Relationship between max(KS) and connections recovered. (**E**) Relationship between max(KS) and the correlation between empirical and model degree. (**F**) Relationship between max(KS) and the correlation between empirical and model degree. (**G-I**) Models using a multiplicative form with exponential distance decay: (**G**) Relationship between max(KS) and connections recovered. (**H**) Relationship between max(KS) and the correlation between empirical and model degree. (**I**) Relationship between max(KS) and the correlation between empirical and model degree. (**J-L**) Models using a multiplicative form with power-law distance decay: (**J**) Relationship between max(KS) and connections recovered. (**K**) Relationship between max(KS) and the correlation between empirical and model degree. (**L**) Relationship between max(KS) and the correlation between empirical and model degree.

**Figure S7.**
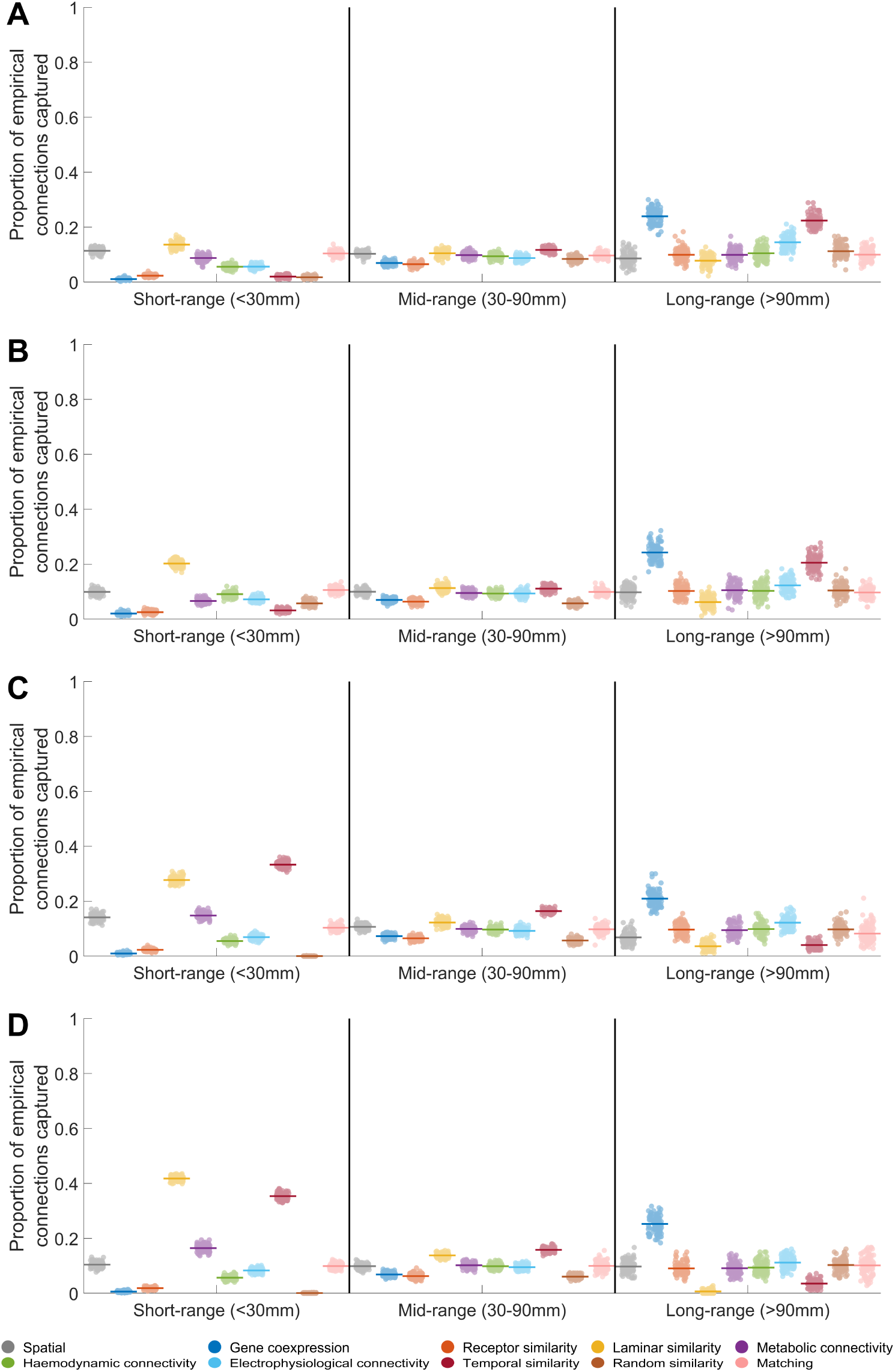
Connections captured at different distance thresholds for generative network models with the strongest degree correlations. Proportion of empirical short-range, mid-range, and long-range connections captured by the generative network models with the strongest degree correlations for the: (A) additive and exponential distance decay formulation; (B) additive and power-law distance decay formulation; (C) multiplicative and exponential distance decay formulation; (D) multiplicative and power-law distance decay formulation. The coloured line indicates the average overlap, while each point indicates the result for an individual model network.

**Figure S8.**
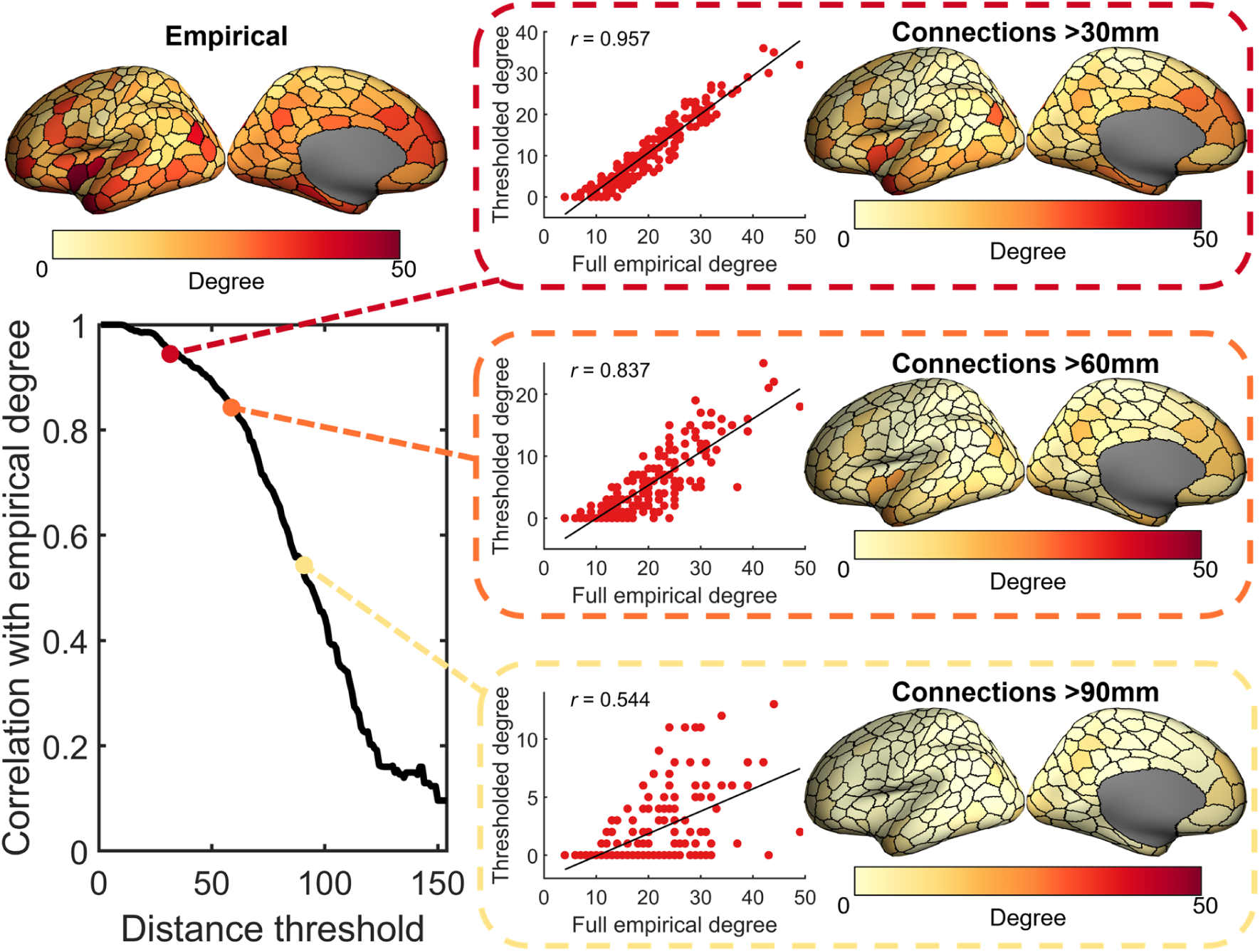
Similarity of the empirical degree sequence at different distance thresholds. Degree is calculated only using connections with a length greater than the current distance threshold. The thresholded degree is then correlated with the full empirical degree (i.e., degree calculated when using all empirical connections).

**Figure S9.**
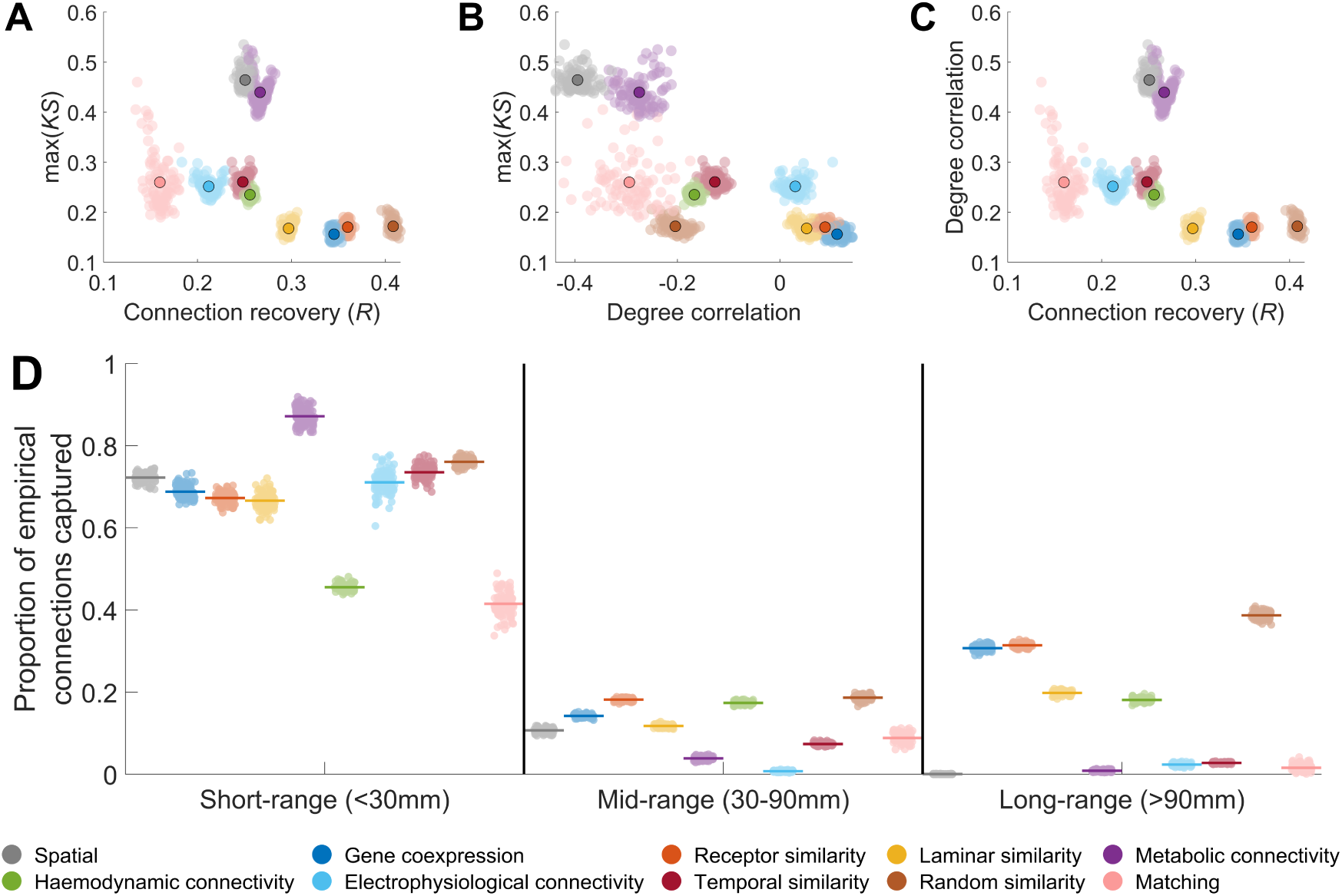
Performance of generative network models in capturing topological and topographical properties in the whole-brain group consensus network. **(A)** Relationship between max(KS) and connection recovery of synthetic model networks. **(B)** Relationship between max(KS) and the correlation between empirical and model node degree. **(C)** Relationship between connection recovery and the correlation between empirical and model degree. In each plot, the outlined point indicates the average across model runs for different model formulations. (**D**) Proportion of empirical short-range, mid-range, and long-range connections captured by the best fitting generative network models. The coloured line indicates the average overlap, while each point indicates the result for an individual model network.

**Figure S10.**
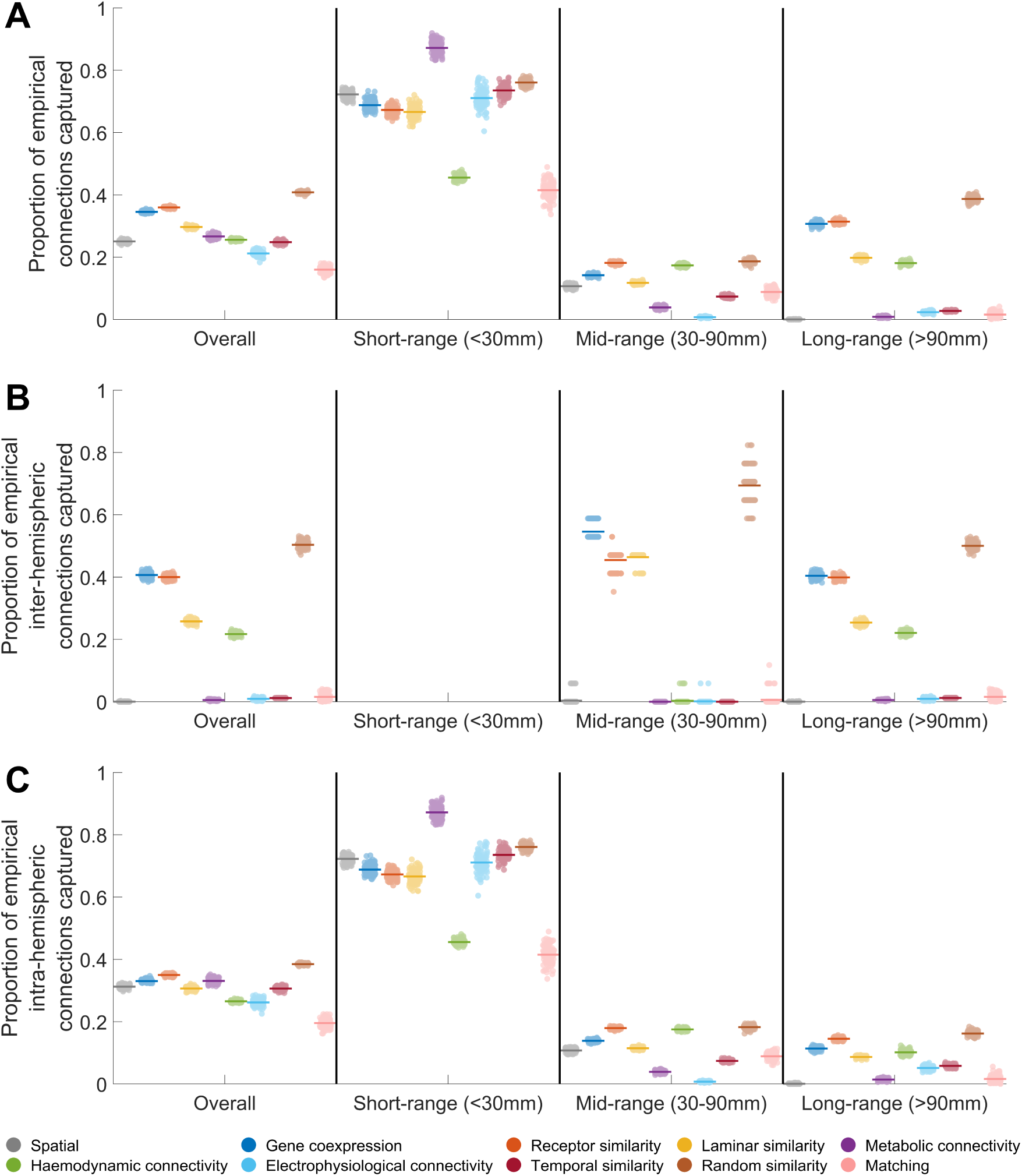
Connection recovery for all, inter-hemispheric, and intra-hemispheric connections for the whole-brain group consensus network by the additive, exponential decay GNMs. Proportion of empirical connections captured for (A) all connection types; (B) inter-hemispherical connections; (C) intra-hemispheric connections across different distance thresholds (all, short-range, mid-range, and long-range). Note that no short-range inter-hemispheric connections exist, and the stratification for the mid-range inter-hemispheric connections is because few such connections exist empirically.

**Figure S11.**
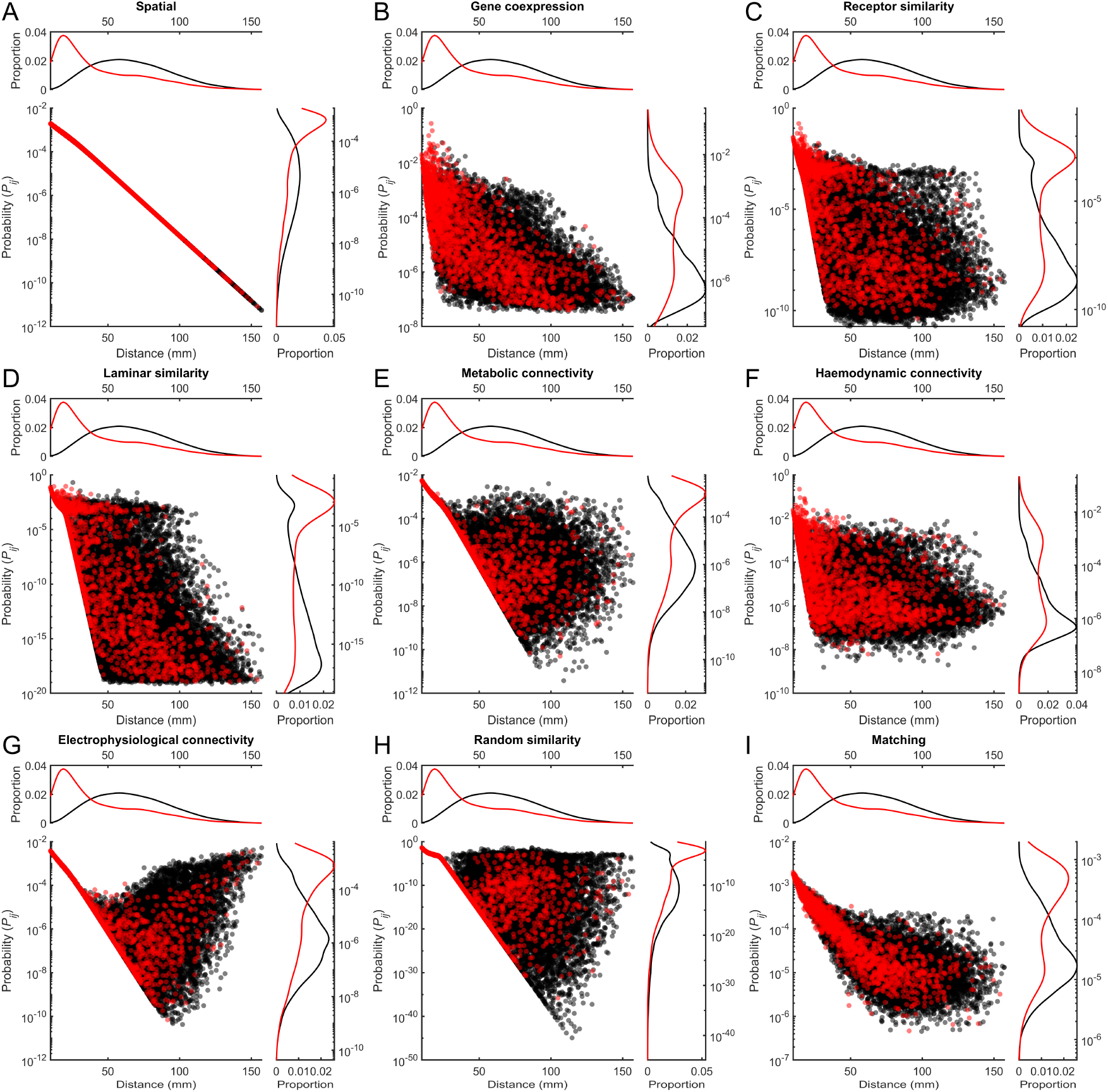
Connection probabilities in generative network models. Shown are connection probabilities for the network with the lowest max(KS) for all GNMs besides temporal similarity (see figure in main text). The connection probabilities are the mean probability assigned to each connection across all timepoints of the model. Red points indicate structural connections (i.e., connections that exist in the empirical data) while black points indicate non-structurally connected regions. Kernal density plots for the structural and non-structural connections are shown for the distributions of distance/length and probability against the respective axis.

**Figure S12.**
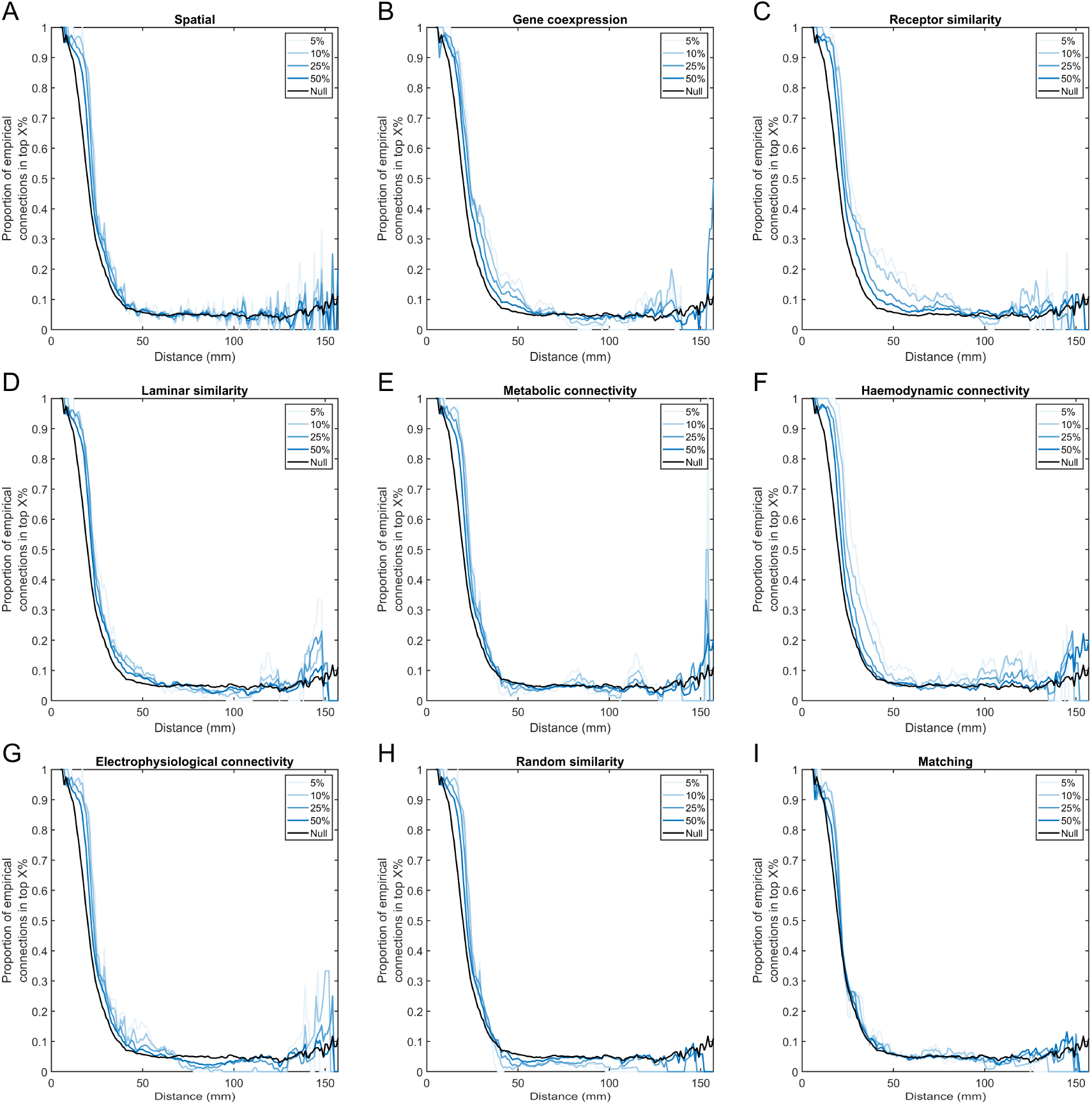
Most probable connections at different distances across generative network models. Using the sliding window analysis, at each distance the top X% of connections with the highest probability are found. The proportion of these which are empirical connections is then calculated. The “null” line indicates the proportion of empirical structural connections that exist at that distance (i.e., the probability of selecting an empirical structural connection if all connections at a given distance were equally probable).

**Figure S13.**
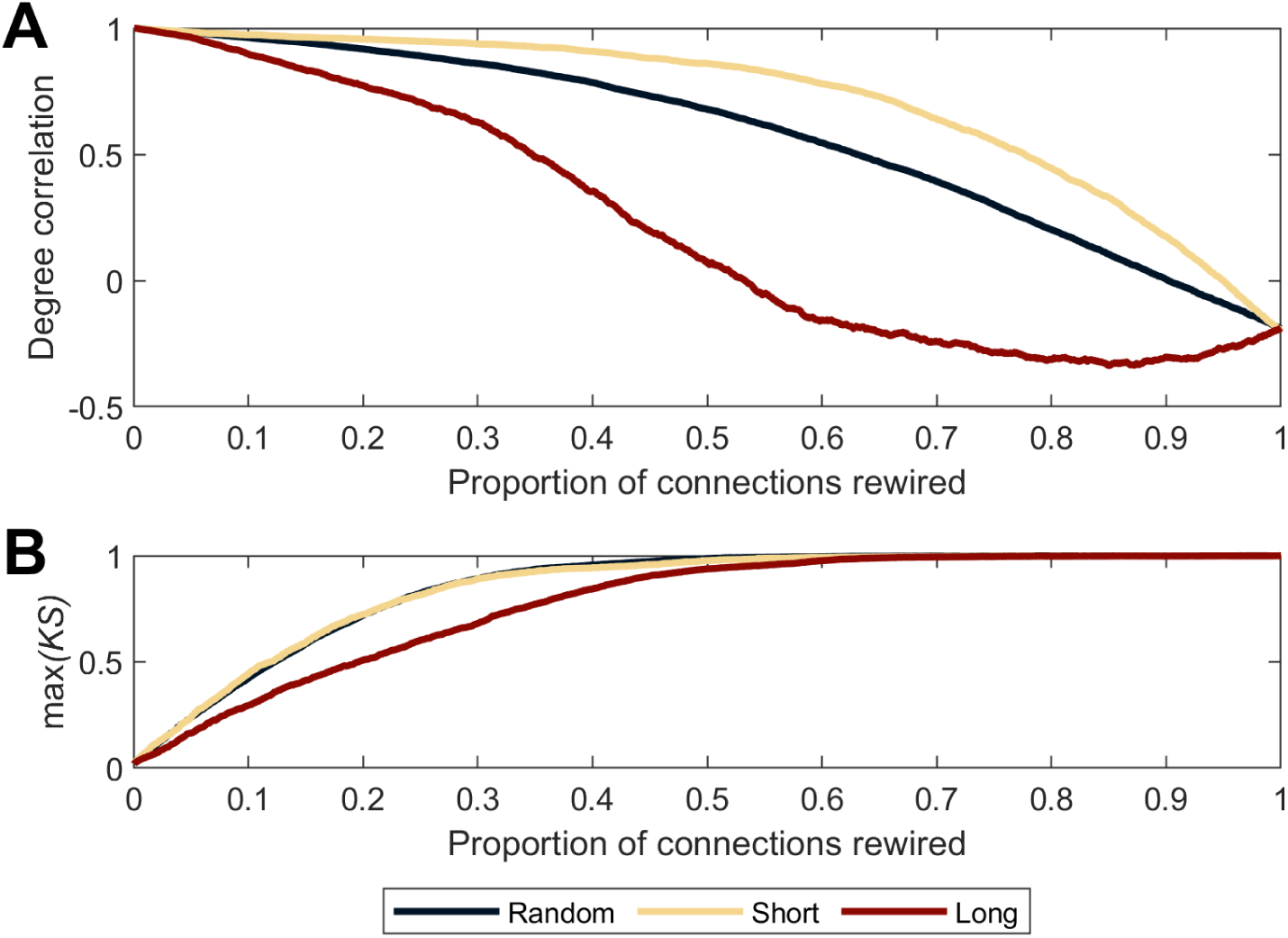
Effect of iteratively rewiring connections on the degree correlation. (A) The proportion of connections rewired and its effect on the degree correlation (i.e., correlation between node degree in the original and rewired network) for different rewiring algorithms. (B) The proportion of connections rewired and its effect on max(KS) for different rewiring algorithms. The *random* algorithm rewired connections at random but of a similar length; *short* rewired connections from shortest to longest connections with those of a similar length and; *long* rewired connections from longest to shortest connections with those of a similar length.

**Figure S14.**
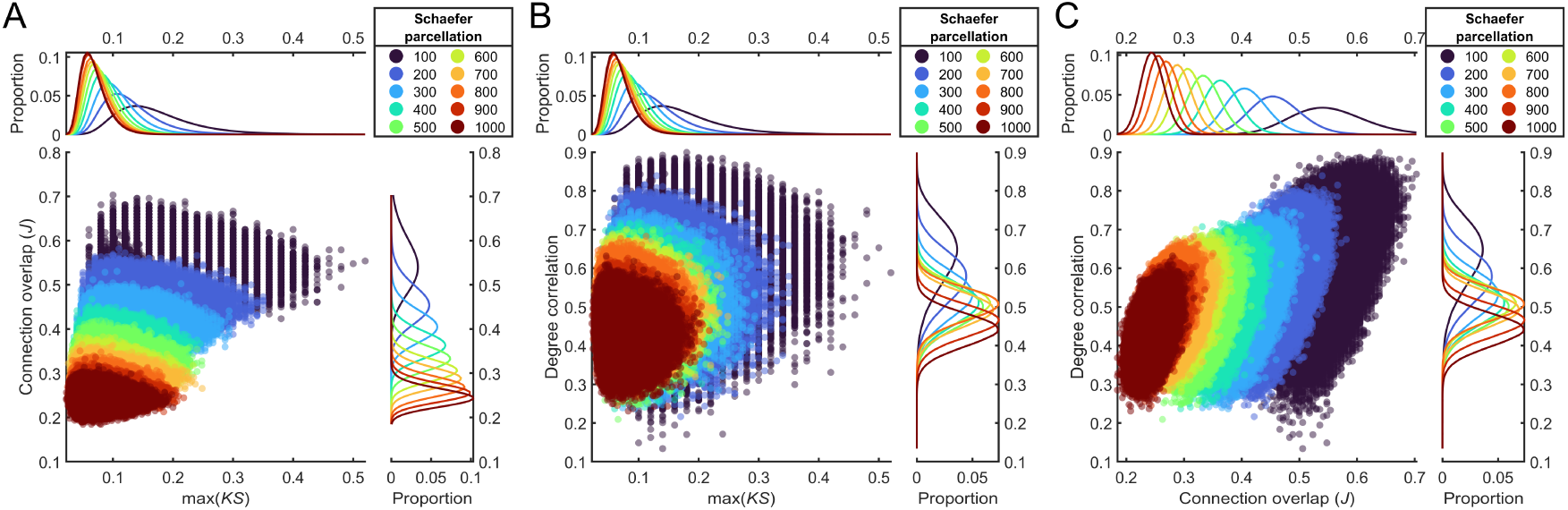
Performance of max(KS), connection overlap, and similarity of the degree distribution when comparing empirical networks produced using deterministic tractography. **(A)** Relationship between max(KS) and connection overlap. **(B)** Relationship between max(KS) and the correlation between two empirical networks nodal degree. **(C)** Relationship between connection overlap and the correlation between two empirical networks nodal degree. The kernel density plots show the distributions for each parcellation on the respective feature.

**Figure S15.**
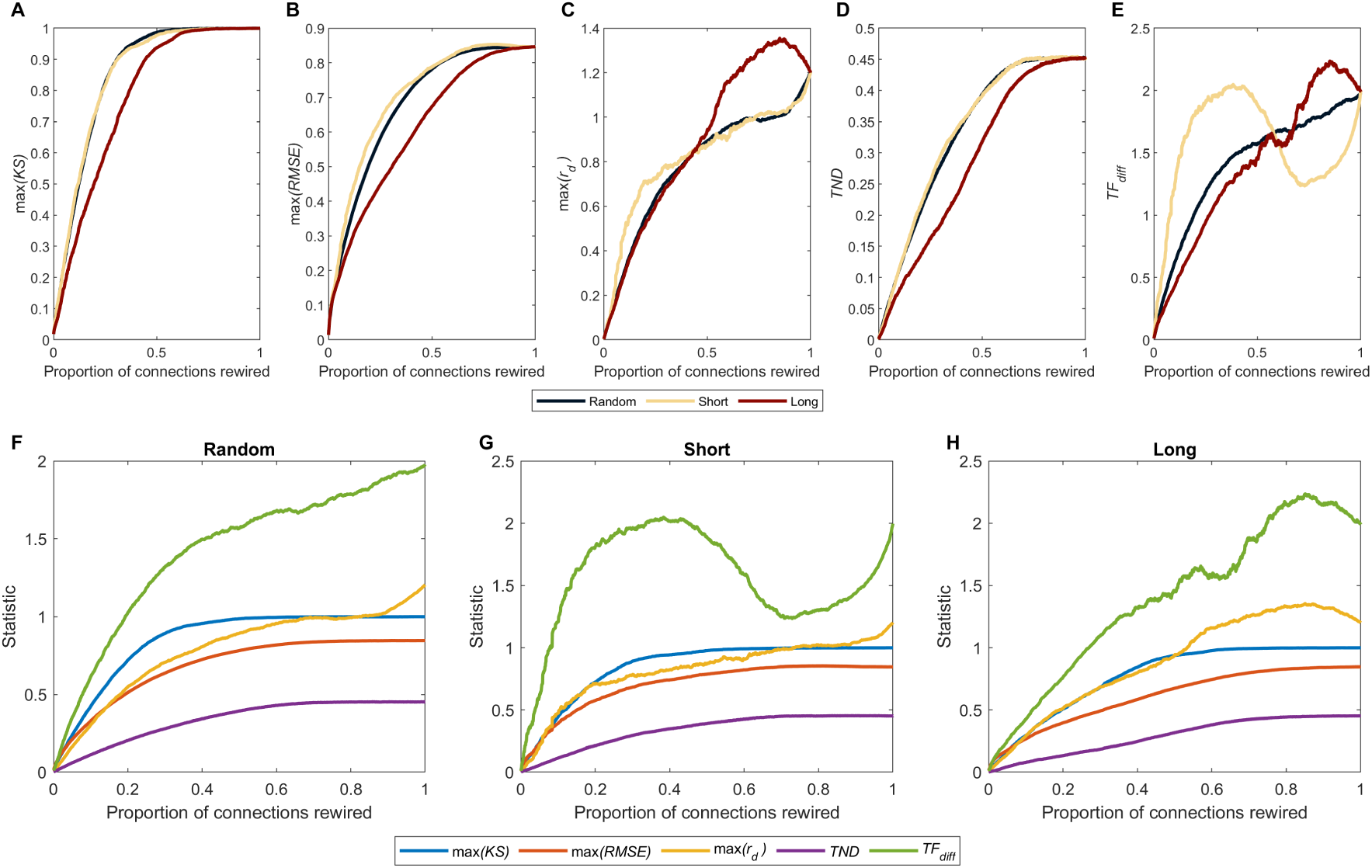
Effect of iteratively rewiring edges on different network similarity measures. The proportion of connections rewired and its effect on (A) max(KS); (B) max(RMSE); (C) max(r_d_); (D) TND; (E) and TF_diff_ for different rewiring algorithms. Each statistic has been normalised to its value obtained at 100% rewiring. Comparison of the different similarity measures for the: (F) *random*: (G) *short*; (H) *long*. The random algorithm rewired connections at random but of a similar length; *short* rewired connections from shortest to longest connections with those of a similar length; *long* rewired connections from longest to shortest connections with those of a similar length.

---

**Table S1.**
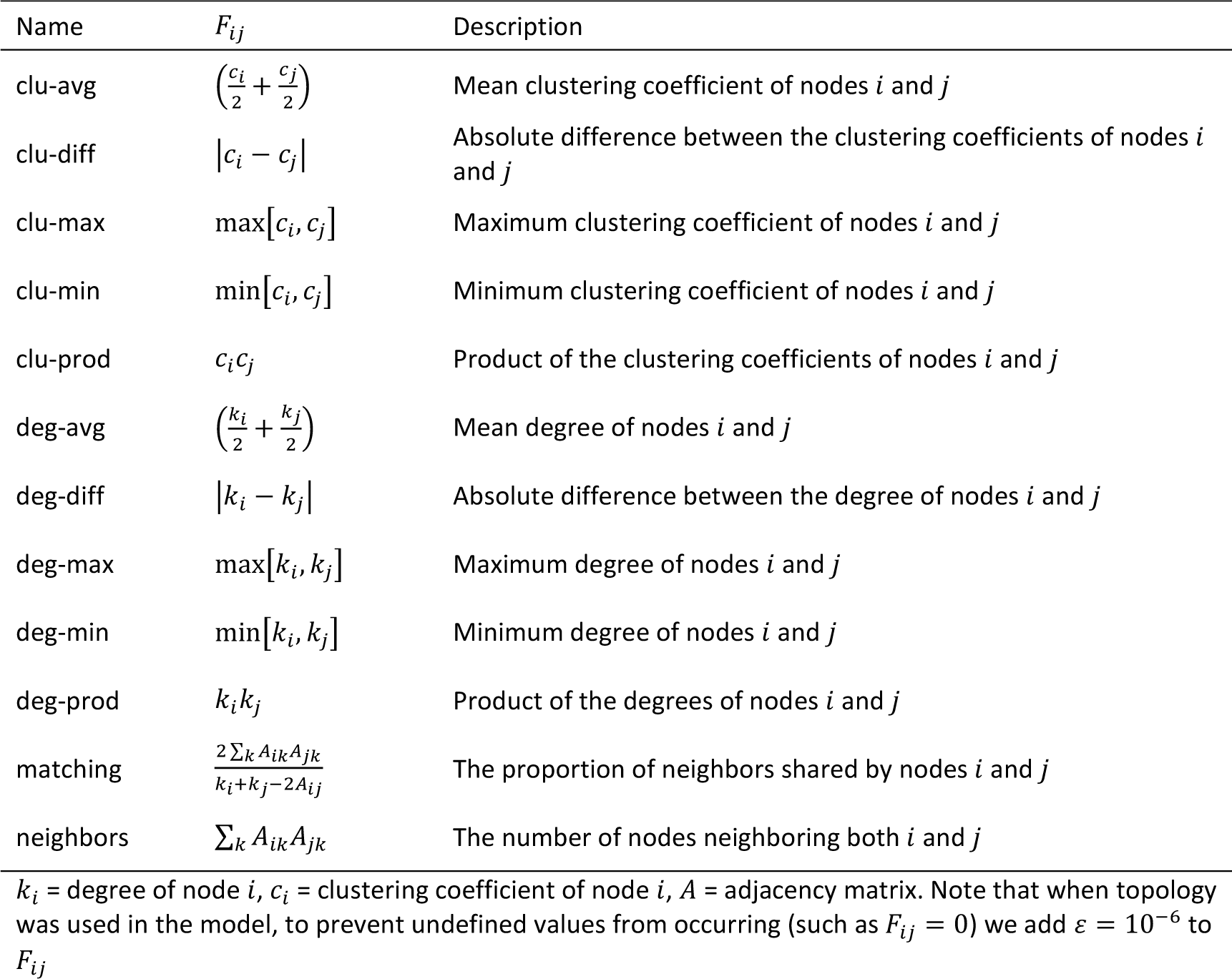
Definitions of topological coupling terms.

**Table S2.**
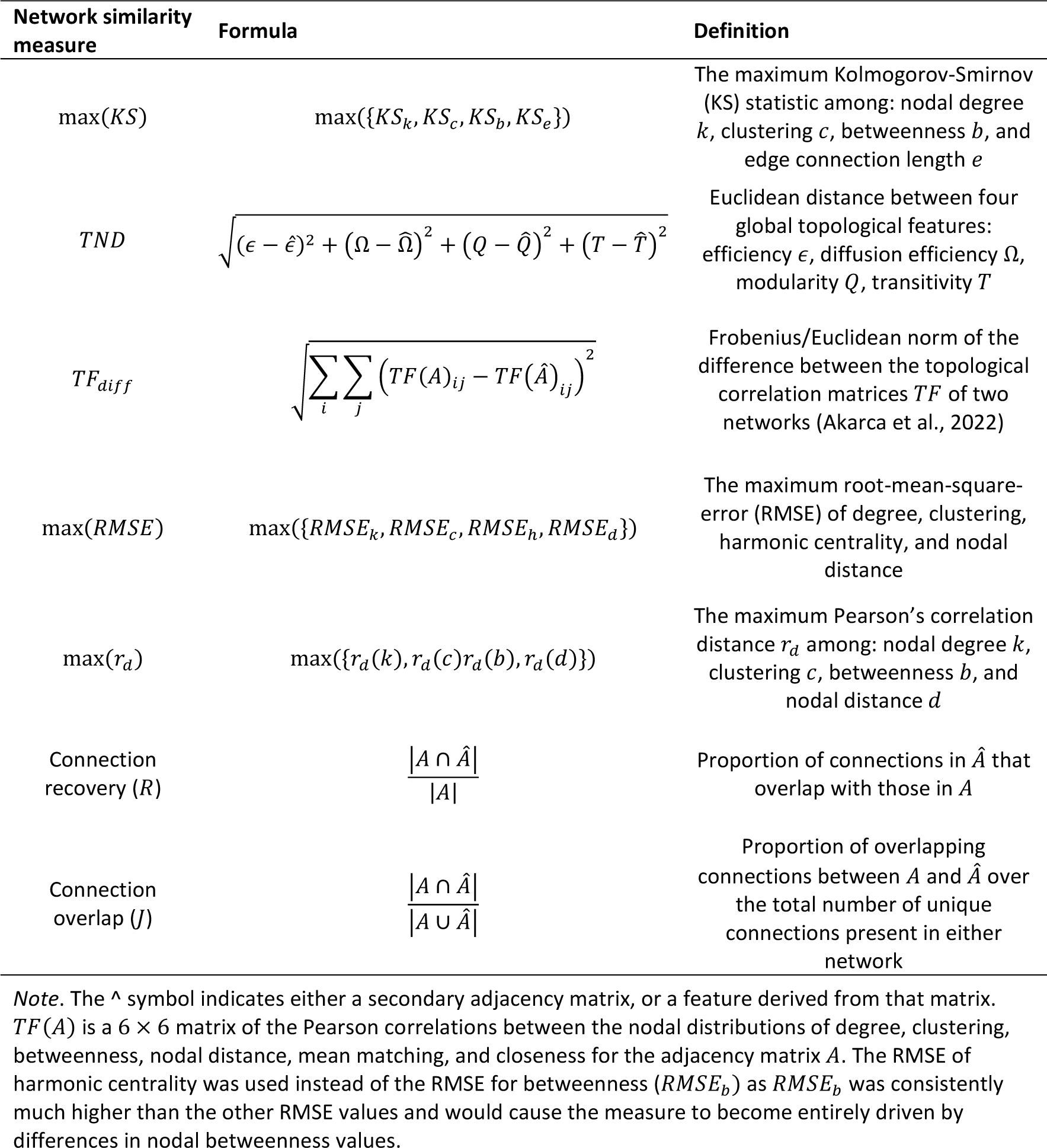
Different measures of network topological and topographical similarity.

**Table S3.**
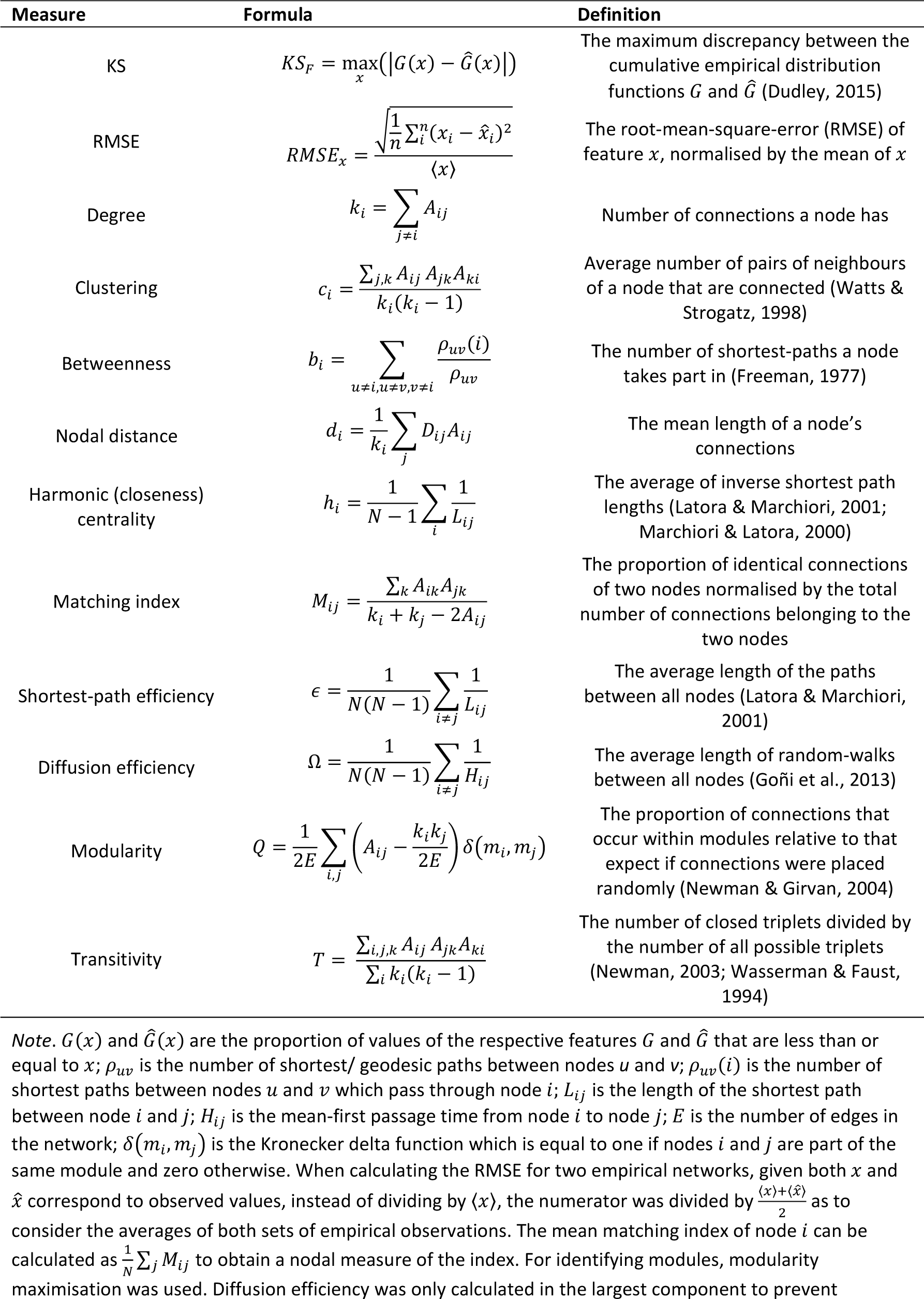
Network measures/miscellaneous equations.

